# Cluster nanoarchitecture and structural diversity of PIEZO1 at rest and during activation in intact cells

**DOI:** 10.1101/2024.11.26.625366

**Authors:** Clement Verkest, Lucas Roettger, Nadja Zeitzschel, Stefan G. Lechner

## Abstract

The force-gated ion channel PIEZO1 confers mechanosensitivity to many cell types. While the structure and physiological roles of PIEZO1 are well-described, the subcellular distribution and the impact of the cellular microenvironment on PIEZO1 conformation and function are poorly understood. Here, using MINFLUX nanoscopy we demonstrate that PIEZO1 channels collectively deform the membrane into pit-shaped invaginations, thereby creating mechano-responsive microdomains capable of amplifying mechanical stimuli via subtle changes in their topology. Moreover, by measuring intramolecular distances in individual PIEZO1 channels with nanometer precision, we reveal subcellular compartment-specific differences in PIEZO1 conformation at rest and during activation that correlate with differences in PIEZO1 function and are possibly caused by differences in cytoskeletal architecture. Together, our data provide previously unrecognized insights into the complex interplay of forces that determine how PIEZO1 alters membrane shape and, vice versa, how the membrane together with the cytoskeleton affect the conformation and function of individual PIEZO1 channels.

**Teaser:** MINFLUX nanoscopy reveals subcellular distribution and conformational diversity of PIEZO1 channels in intact cells.

## Introduction

PIEZO1 is a mechanosensitive ion channel that enables a variety of cell types to detect and respond to mechanical stimuli and is thus essential for numerous physiological processes (*1–5*). With regards to PIEZO1 structure and function, cryogenic electron microscopy (cryo-EM) studies together with high-speed AFM and in-silico modelling have established a mechanistic framework, which proposes that PIEZO1 forms a propeller-shaped homotrimer that assumes a curved conformation at rest and transitions – via a partially flattened and intermediate open state – into a fully flattened conformation (*6–17*) upon activation. In the flattened state, the channel can be ‘open’ or ‘inactivated’, depending on whether the cap-gated is closed or not (*14*).

This framework, however, rests on investigations of individual channels in artificial membranes and simplified model systems and thus has several limitations. Firstly, in cryo-EM liposomes, PIEZO1 adopts different conformations depending on its orientation: Thus, in concave environments (i.e. when reconstituted in the inside-out orientation), PIEZO1 assumes a curved conformation (*15–17*), whereas it assumes the flattened conformation when reconstituted in the outside-out orientation (*7*) (Fig. 1A). Cryo-EM liposomes are, however, highly curved and only slightly bigger that PIEZO1 itself, such that the channel is exposed to abnormally high bending forces when reconstituted in such liposomes, which raises the question if the conformational states resolved by cryo-EM also exist in the native environment of the plasma membrane where such high degrees of membrane curvature are not observed. Secondly, in living cells the membrane and possibly the channel itself, are tightly attached to the subjacent cytoskeleton, which is known to control PIEZO1 sensitivity (*18–22*). Yet, it is unclear if and to what extent the cytoskeleton alters PIEZO1 conformation and changes thereof upon mechanical activation. Mulhall et al. recently reported that PIEZO1 appears to be more expanded in living cells than predicted by cryo-EM, indicating that the channel indeed behaves differently in its native environment (*23*), though the cell intrinsic factors causing the observed partial flattening remained elusive.

**Fig. 1.**
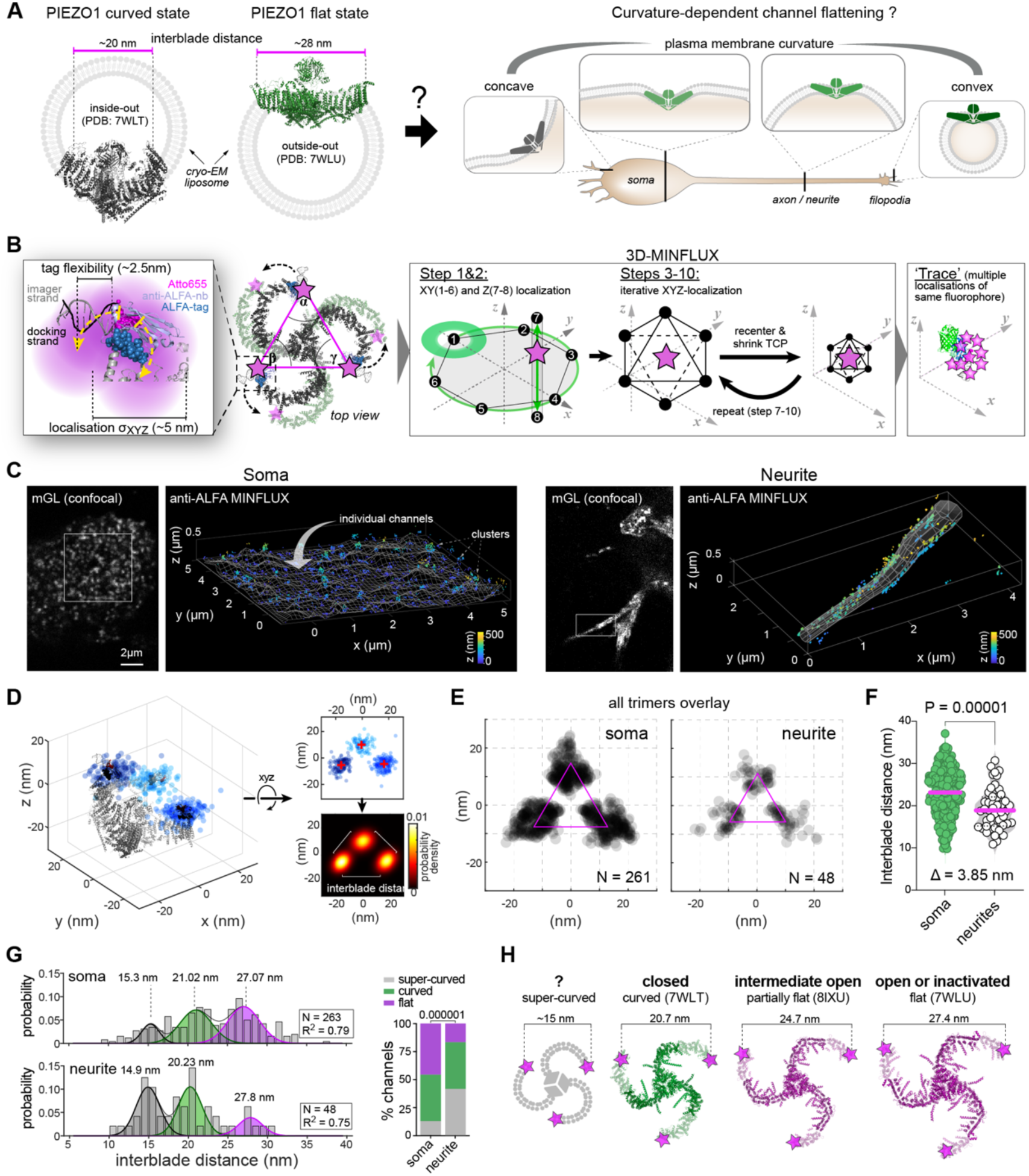
3D-MINFLUX reveals cell compartment-specific conformational heterogeneity of PIEZO1. **(A)** Cartoon representing the side views of curved (black) and flattened (green) PIEZO1 conformations determined by cryo-EM in liposomes (left), together with the possible source of curvature variation in living cells (right). **(B)** Close-up view on the peripheral extracellular blade of PIEZO1-ALFA-mGL together with the ALFA nanobody to illustrate linkage and localization precision during MINFLUX DNA-PAINT imaging. Distances between ALFA-tags are further depicted (pink) as well as the angles between the lines connecting the ALFA-tags (left), along with a cartoon illustrating the principle of 3D-MINFLUX/DNA-PAINT (right). **(C)** confocal images (left) localizations of a soma and a highly curved neurite showing PIEZO1-ALFA-mGL expression and perspective view of the corresponding MINFLUX localization data (right) from the marked regions. **(D)** 3D scatter plots of raw localizations of a representative trimer (left), with superimposed cryo-EM structure, found in a cell soma. The 2D in-plane projections of the 3D data were fitted with a bivariate Gaussian distribution, with their probability densities (right). **(E)** Super-particle of all trimers found in somata (left) or neurite (right) generated by aligning the trace means of each identified trimer to a reference trimer. **(F)** Violin plots of interblade distances of trimers residing in somata (green, N=261 trimers from 15 cells) and neurites (grey, N=48 trimers from 15 cells). Comparison using Mann Whitney test. **(G)** Frequency distribution (left) of interblade distances measured in soma (top) and neurite (bottom). Distribution was fitted with a Gaussian mixture model. The three components are highlighted in grey, green and pink, with their peak values and the goodness of the fit (R²). Proportion of the three observed PIEZO1 classes (right) in soma and neurite, determined with the fitted values. Comparison with Chi-Square test. **(H)** Structural models of PIEZO1 from cryo-EM. Peripheral parts of the blades missing from cryo-EM data were modelled using the AlphaFold structure (transparent).

Finally, although it is well-established that individual PIEZO1 channels can function as independent mechanosensors (*24*, *25*), PIEZO1 was shown to form prominent clusters in many cell types, both at endogenous expression levels and when overexpressed in heterologous systems (*20*, *26–34*). Using immunohistochemistry and genetic fluorophore tagging of PIEZO1, prominent cluster formation at endogenous expression levels was observed in rodent and human tissues including keratinocytes (*33*, *34*), endothelial cells (*34*), neural stem cells (*34*), dermal fibroblasts (*35*), glioblastoma stem cells (*26*), microglia (*29*), cardiomyocytes (*32*) and trigeminal ganglia (*31*). Interestingly, PIEZO1 does not appear to form clusters in red blood cells, as suggested by STED super-resolution microscopy (*36*), though conflicting results were obtained using force-distance based atomic force microscopy (*37*). Regarding the functional relevance of PIEZO1 clusters, two elegant studies in which calcium sensors were fused to the C-terminus of PIEZO1, demonstrated that clusters mediate calcium influx in response to mechanical and chemical activation of PIEZO1 (*34*, *38*). Moreover, in some cell types PIEZO1 clusters are recruited to hot spots of mechanotransduction such as focal adhesions (*26*, *28*, *30*), the rear edge of migrating keratinocytes (*33*) and t-tubules in cardiomyocytes (*32*), which further supports an important role of PIEZO1 clusters in cellular mechanosensitivity. In the curved conformation PIEZO1 locally deforms the membrane into a dome shape that extends far beyond the perimeter of the channel, such that the membrane footprint of PIEZO1 is much bigger than the channel itself (*15–17*) (Fig. 1A), which supposedly has important implications regarding the impact of clustering on PIEZO1 conformation and function. Thus, it has been proposed that the opposing curvatures of the membrane footprints of two nearby channels would create an energetic constraint in the interjacent membrane such that nearby channels would either repel each other or induce mutual flattening to reduce the overall energy of the system (*6*, *12*, *39*). These hypotheses have, however, never been directly tested. Using 2D stochastic optical reconstruction microscopy (STORM), Ridone and colleagues demonstrated that PIEZO1 clusters are densely packed and contain multiple channels, but due to the lack of 3D information the exact number of channels per cluster and their spatial arrangement remained elusive (*27*).

Here, we thus set out to examine the nanoarchitecture and the distribution of PIEZO1 within clusters as well as the possible role of other cell intrinsic factors in controlling PIEZO1 conformation at rest, using 3D-MINFLUX nanoscopy in combination with DNA-PAINT (*40–42*).

## Results

### PIEZO1 conformation at rest differs between subcellular compartments

In proteoliposomes used for cryo-EM, PIEZO1 assumes a curved conformation with upward-tilted blades when reconstituted in the inside-out orientation (*15–17*), whereas it assumes a flattened conformation in which the distance between the distal ends of the blades is increased by ∼8 nm (*7*) when reconstituted in the outside-out orientation (Fig. 1A). Accordingly, it has been proposed that membrane curvature governs PIEZO1 conformation, which raises the question as to whether cell compartment-specific differences in membrane curvature are sufficient to fine-tune PIEZO1 conformation and hence function at rest (Fig. 1A). To test this hypothesis, we measured the interblade distances of PIEZO1 trimers located at the flat soma-substrate interface and in neurites with highly curved membranes, using MINFLUX nanoscopy in combination with DNA-PAINT, which achieves spatial resolutions in the nanometer range (Fig. 1B, (*40–42*)). To this end, we expressed a PIEZO1 variant that carries an extracellular ALFA-tag at the distal end of the blade domain and an mGreenLantern-tag in the C-terminus (PIEZO1-ALFA-mGL, Fig. S1) in Neuro2a-PIEZO1 knock-out cells (*43*), labelled the ALFA-tags with an anti-ALFA single domain nanobody conjugated to a DNA-PAINT docking-strand and imaged the cells with an Atto655-conjugated DNA-PAINT imager strand using 3D-MINFLUX (Fig. 1C and Fig. S2 for negative controls and raw data processing).

To identify triple-labelled PIEZO1 trimers in MINFLUX localisation data, we computed the Euclidian distance matrix of all localisations and searched for localisation triplets in which the distances between the individual localisations were smaller than 40 nm (i.e. the maximal physically possible distance between two ALFA-tags in the fully flattened state) and that had no other neighbouring signals within 60 nm (see Fig. S3A and B as well as methods for details on trimer identification rules). Using this algorithm, we identified 261 trimers in somata and 48 in neurites (Fig. 1C-E, Fig. S4A and Supplementary Video S1 and 2), with localization errors around 4 nm in *xyz* (Fig. S3C). Note, due to the stringent trimer identification criteria no trimers were detected in densely packed PIEZO1 clusters (Fig. 1C). To compare overall trimer geometries, we aligned the localization trace means of each trimer to a 2D reference trimer using an iterative closest point algorithm and overlaid the data. These overlays revealed elongated localization clouds extending outward from the trimer center, indicating that variability does not only result from the isotropic MINFLUX localization error (Fig. 1E), but also reflects varying degrees of PIEZO1 flattening. Moreover, interblade angles (i.e. angles between the lines connecting the distal ends of the blades; Fig. 1B), while being uniformly distributed around 60°, showed notable variability (Fig. S4B), indicating additional rotational flexibility of the distal blades, as previously reported (*23*).

Most notably, we found that the mean interblade distance was significantly smaller in neurites than in somata (neurite: 19.3 ± 4.9 nm; soma: 23.1 ± 5.6 nm; P = 0.00001, Mann– Whitney test; Fig. 1F), indicating that PIEZO1 is generally more contracted in neurites. Analysis of interblade distance distributions using Gaussian mixture models further revealed three distinct conformational states with interblade distance means matching previously described cryo-EM structures. Thus, 45.6 % of the trimers detected in somata, adopted a conformation with a mean interblade distance of 27.07 nm, which was similar to the interblade distance measured in the cryo-EM structure of the flattened conformation (PDB:7WLU), and 41.4 % of the channels exhibited a mean interblade distance of 21.02 nm, which was almost identical to the interblade distance of the curved conformation (PDB:7WLT, Fig. 1G and H). Interestingly, we found that a considerable fraction of the channels (13 %) adopts a conformation that is even more contracted than the curved conformation – hereafter referred to as ‘super-curved’ – with a mean interblade distance of 15.3 nm. These same conformations were observed in neurites (flat: 27.8 nm, curved: 20.2 nm, super-curved: 14.9 nm), but the proportions of the three subclasses significantly differed. Thus, while almost half of the channels were flat in the soma, only very few channels (16.7 %) adopted the flat conformation in neurites and, accordingly, significantly more channels were curved and super-curved (Fig. 1G and H).

Hence, our data indicates that PIEZO1 preferentially adopts certain conformational states that closely match those previously described by cryo-EM and shows that the preference for these states differs between subcellular compartments.

### PIEZO1 in neurites is more susceptible to Yoda1-induced flattening

We next examined the implications of the differential preference for specific conformational states in neurites and somata with regards to PIEZO1 function. Assessing PIEZO1 mechanosensitivity in neurites is challenging due to limitations of standard patch-clamp techniques. In cell-attached pressure-clamp recordings, seal creep (*44*) induces F-actin reorganization (*45*), making this approach unsuitable for studying effects linked to cytoskeletal differences between neurites and soma. Whole-cell recordings also face issues, as space-clamp limitations prevent control of neurite membrane potential, precluding meaningful current comparisons. To overcome this, we used jGCamp8 calcium imaging in intact cells to explore PIEZO1 sensitivity. Neurites showed greater responses to 100 nM and 300 nM Yoda1 compared to somata, though responses converged at higher concentrations (Fig. 2A), indicating that PIEZO1 is more sensitive in neurites as compared to somata.

**Fig. 2.**
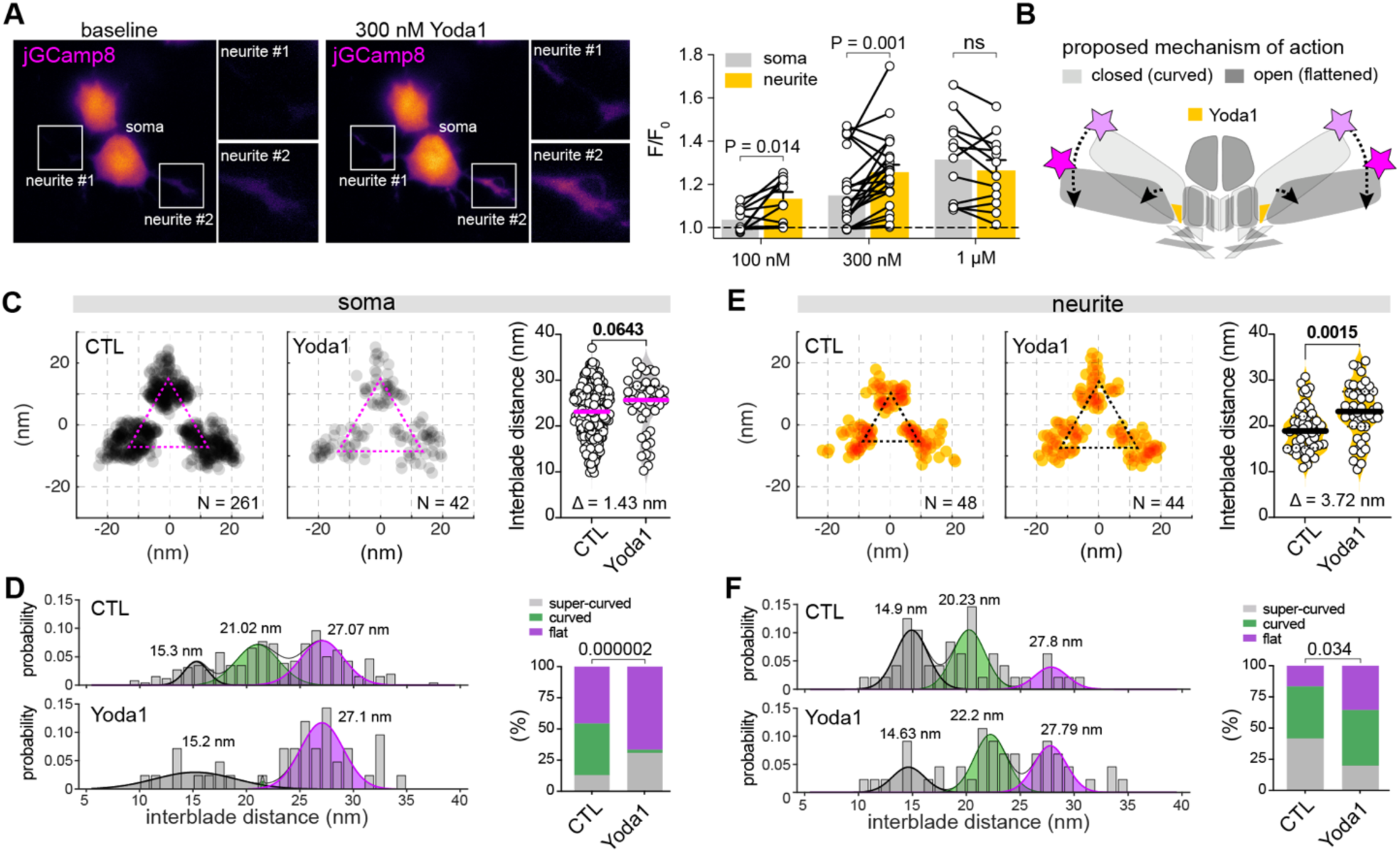
PIEZO1 in neurites is more susceptible to Yoda1-induced flattening. **(A)** Representative wide-field fluorescence images (left) of N2a-P1KO cells coexpressing PIEZO1mScarlet and jGCaMP8m before and after the application of 300nM Yoda1. Insets are close-up view of neurites. Average of the maximal Yoda1 effect (right) on the normalized fluorescence (F/F0) at different concentrations between cell soma and theirs associated neurites. Comparison with paired t-test. N=10 (100nM), N=23 (300nM) and N=12 cells (1µM). **(B)** Cartoon depicting the proposed mechanism of action of Yoda1 on PIEZO1. **(C-F)** Superparticle of all trimers from control and cells treated with Yoda1 in soma (C) and in neurites (E) (left) and comparison of the interblade distances with and without Yoda1 in soma and neurite, with Mann-Whitney test (right). Frequency distribution of interblade distances measured in control or Yoda1 treated cells in soma (D) and in neurites (F). Distribution was fitted with a Gaussian mixture model. The three components are highlighted in grey, green and pink (left). Comparison of the proportion of the three observed PIEZO1 classes with or without Yoda1 in soma and neurites, determined with the fitted values (right). Comparison with Chi-Square test.

Yoda1 is thought to activate PIEZO1 by acting as a molecular wedge that binds to a hydrophobic pocket at the proximal end of the blade domain, thereby promoting channel flattening (Fig. 2B, (*46*)). We thus next asked whether the increased Yoda1 sensitivity in neurites corresponds to greater conformational changes in PIEZO1. Consistent with a prior report (*23*), we observed a subtle – yet non-significant – increase in the average interblade distance in the somata of cells fixed in the presence of Yoda1 (Δ = 1.43 nm, P = 0.0643; Fig. 2C). Gaussian mixture modelling, however, revealed a significant shift toward the flat conformation and an almost complete absence of curved channels (Fig. 2D). The super-curved population also increased, explaining the small overall change in mean interblade distance. In neurites, by contrast, Yoda1 significantly increased mean interblade distance by 3.72 nm (P = 0.0015, Fig. 2E), with more flat and fewer super-curved channels (Fig. 2F). The ‘curved’ population remained constant in proportion but showed an increase in mean interblade distance from 20.23 nm to 22.2 nm, making the overall effect of Yoda1 more pronounced in neurites than somata (Fig. 2F).

Hence, our data reveal differences in Yoda1-induced conformational changes in PIEZO1 between channels located in neurites and somata and suggest that the susceptibility to flattening correlates with its chemosensitivity.

### Local differences in cytoskeletal rigidity govern PIEZO1 conformation

The unexpected observation that PIEZO1 was more contracted in highly curved neurites as compared to the flat soma-substrate interface (Fig. 1), prompted us to explore the role of factors other than membrane curvature, that could differentially alter PIEZO1 conformation in neurites and somata. First, we examined the relationship between interblade distance and neurite diameter within the group of trimers detected in neurites, where curvature varies, but other cell intrinsic factors likely do not. We did, however, not observe any correlation (Fig. 3A), suggesting that membrane curvature – at least at the micro scale – only plays a minor role in controlling PIEZO1 conformation. Another possible explanation for the observed differences in interblade distances comes from modelling studies, which have proposed that crowding of PIEZO1 may induce mutual flattening of adjacent channels (*39*). However, there was no correlation between local channel density – i.e. average distances to the three nearest neighbouring channels of the detected trimers – and the interblade distance within the two groups (Fig. 3B), indicating that differences in crowding do not contribute to the observed differences in interblade distances. A possible confounding factor of our data is the imaging depth of MINFLUX, which is limited to a few hundred nanometers, such that the apical surface of the cell soma cannot be imaged. Accordingly, all trimers that were detected in somata resided at the cell-substrate interface, where the channels could possibly be tethered to the extracellular matrix (ECM), whereas in neurites trimers were detected in the basement membrane as well as in the apical membrane. There was, however, no difference between the interblade distances of trimers in the basement membrane of neurites compared to those residing in the apical membrane, indicating that possible tethering of PIEZO1 to the ECM or lack thereof in the apical membrane does not alter PIEZO1 conformation (Fig. 3C).

**Fig. 3.**
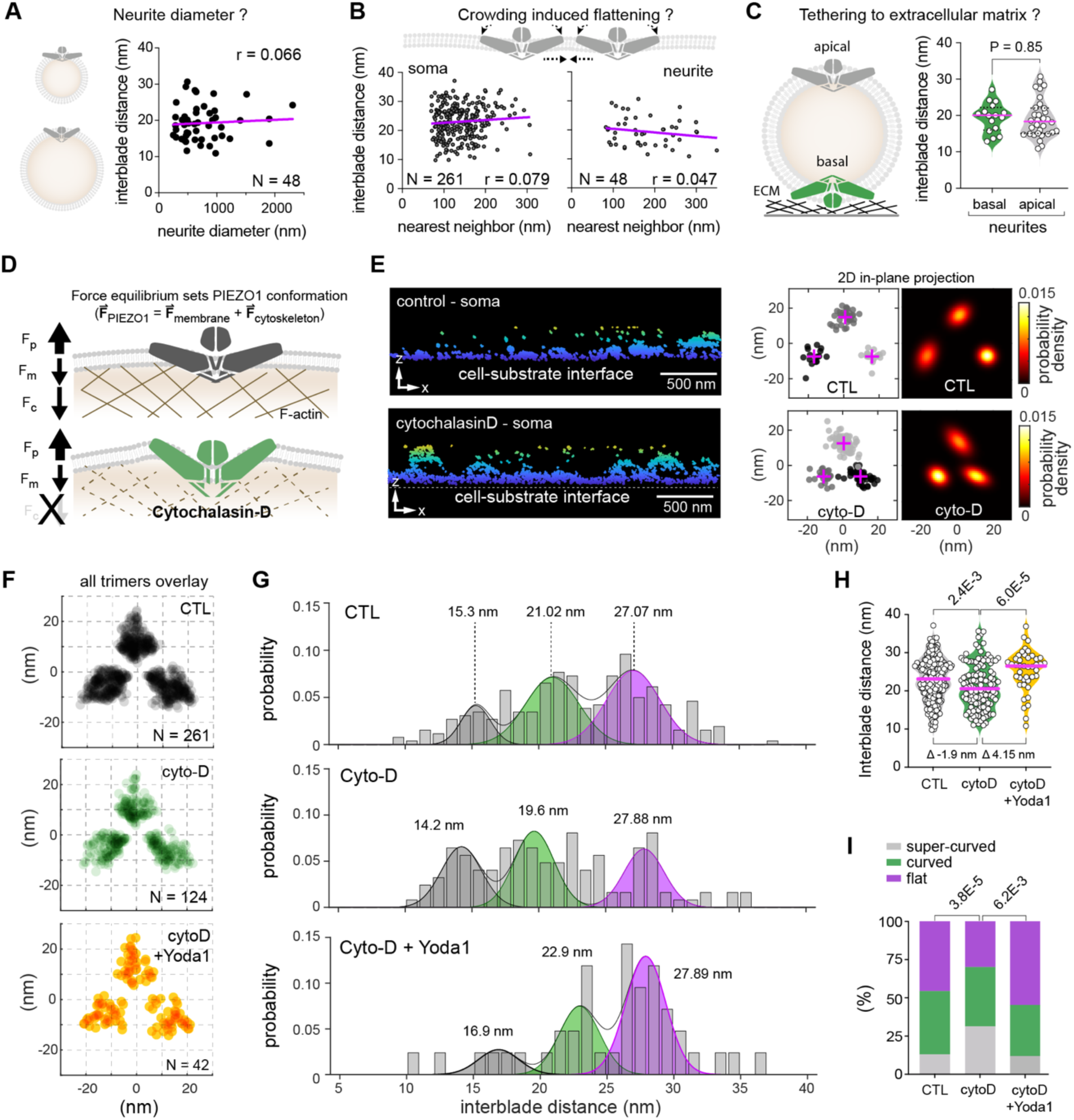
Disruption of the cytoskeleton alters PIEZO1 conformation. **(A)** Cartoon highlighting the possible influence of the neurite diameter on PIEZO1 conformation (left) and correlation between measured interblade distance and estimated neurite diameter at PIEZO trimer localization. Pearson correlation coefficient r is indicated. Linear regression is in pink. **(B)** Cartoon illustrating the possible effect of PIEZO1 crowding (top) and correlation between measured interblade distance in the soma and neurites with the average distance of the three nearest neighbours of identified PIEZO1 trimers (bottom). Pearson correlation coefficient r is indicated and linear regression is in pink. **(C)** Cartoon illustrating the possible effect of the ECM in neurites (left) and comparison of the interblade distances between trimer located at the basal or apical side of neurites. Comparison with Mann-Whitney test. **(D)** Cartoon showing the possible interplay of forces exerted by and acting on PIEZO1. **(E)** side views of MINFLUX raw localization data (left) from control (top) and cyto-D-treated cell (bottom), showing that cyto-D does not change the curvature of the cell-substrate interface. 2-D in-plane projections of the raw localization data and heatmaps of the localization probability densities (right) of representative trimer from control and cyto-D treated cells (right). **(F)** Superparticle of all trimers from control (top), cyto-D (middle) and cyto-D + Yoda1 (bottom) treated cells. **(G)** Frequency distribution of interblade distances measured in control (top), cyto-D (middle) and cyto-D + Yoda1 (bottom) treated cells. Distribution was fitted with a Gaussian mixture model. The three components are highlighted. **(H)** Mean interblade distances of all trimers from control (N=15 cells), cyto-D (N=11 cells) and cyto-D+yoda1 (N=3 cells). Comparison with Mann-Whitney tests. **(I)** Proportion of the three observed PIEZO1 classes (right) in control, cyto-D and cyto-D + yoda1, determined with the fitted values. Comparison with Chi-Square tests.

Finally, we considered the role of the cytoskeleton. The plasma membrane is tightly attached to the cytoskeleton, suggesting that the extent to which PIEZO1 deforms the membrane into a dome shape – and thus its own conformation at rest – may be controlled by the equilibrium of forces that PIEZO1 exerts on the membrane and the opposing forces that the membrane together with the cytoskeleton exert on PIEZO1 (Fig. 3D). Since neurites have a different overall cytoskeletal architecture with fewer membrane-cytoskeleton attachments (*47–50*) and PIEZO1 appears more contracted there (Fig. 1), we hypothesized that differences in local membrane deformability between neurites and somata contribute to the observed differences in PIEZO1 conformation in the two compartments. To test this hypothesis, we treated cells with the actin polymerization inhibitor cytochalasin-D (2µM cyto-D), which removes ‘mechanoprotection’ and hence softens the membrane (*19*, *51*). As previously reported (*19*), this increased PIEZO1 current amplitudes and shifted the pressure-response curve toward less negative pressures (CTL: −40.5 ± 11.3 mmHg vs. cyto-D: −29.1 ± 10.5 mmHg, Fig. S5A-C). Notably, cyto-D treatment significantly reduced the mean interblade distance of somatic PIEZO1 trimers (CTL: 23.1 ± 5.6 nm vs. cyto-D: 21.2 ± 5.9 nm) and altered their conformational distribution, with fewer channels in the flattened and more channels in the super-curved state, mirroring the distribution in neurites (Fig. 2F-I). Considering the similarity of PIEZO1 interblade distance distributions in neurites and cyto-D-treated somata (compare Fig. 1G and Fig. 3G) together with the fact that PIEZO1 in neurites was more sensitive to Yoda1 (Fig. 2A) and that cyto-D treatment increased the sensitivity of somatic PIEZO1 (Fig. S5), we next asked if Yoda1-induced channel flattening was also more pronounced after removal of mechanoprotection. Indeed, in cyto-D-treated cells, Yoda1 also significantly increased the mean interblade distance by over 4 nm (CytoD: 21.2 ± 5.9 nm vs CytoD+Yoda1: 25.35 ± 5.43 nm, P=0.0000601, Mann-Whitney test; Fig. 3H), increased the proportion of flattened channels, while reducing the number of super-curved channels, and – notably – shifted the mean interblade distance of channels classified as ‘curved’ to larger values (22.9 nm, Fig. 3G).

Taken together, our data suggest that reduced cytoskeletal rigidity or fewer membrane attachments may underlie the conformational and possibly functional differences of PIEZO1 between neurites and somata.

### PIEZO1 forms clusters that mediate local calcium influx

The data described so far exclusively focused on PIEZO1 channels that resided in isolation. As evident from anti-ALFA MINFLUX scans (Fig. 1C, 3E and 4A) as well as from confocal scans of mGL fluorescence (Fig. 4A), and as previously reported (*20*, *26*, *27*, *29–35*, *52*), however, PIEZO1 channels are not uniformly distributed across the cell surface, but also form prominent clusters that appear to be densely packed with multiple channels. To corroborate previous reports regarding cluster formation, we first examined the distribution of endogenously expressed PIEZO1 in mouse embryonic fibroblasts (MEFs) and U87 human glioblastoma cells using immunocytochemistry and TIRF microscopy. In both cell lines, immunofluorescence signals were scattered across the plasma membrane and appeared as discrete puncta with mean diameters of 362.8 ± 81.3 nm (MEFs) and 375.5 ± 62.2 nm (U87, Fig. 4B), indicating that PIEZO1 also forms clusters at endogenous expression levels that were indistinguishable with regards to size from the clusters observed in N2a cells overexpressing PIEZO1-ALFA-mGL (406.2 ± 97.9 nm, Fig. 4A). Immunolabelling of unpermeabilized N2a cells expressing PIEZO1-ALFA-mGL with a nanobody directed against the extracellularly located ALFA-tag further demonstrated that 76.5 ± 12.7 % of the mGL-positive clusters were embedded in the plasma membrane, whereas only a small proportion appeared to be intracellular vesicles (Fig. 4C-E). To examine the functional relevance of PIEZO1 clusters, we generated a PIEZO1-jGCamp8m fusion protein in which the genetically encoded calcium sensor jGCamp8m was located adjacent to the intracellular exit of the ion conduction pathway, such that local PIEZO1-mediated calcium influx can be detected (Fig. 4F and Fig. S6). Consistent with previous reports using similar approaches (*34*, *38*), stimulation of PIEZO1-jGCamp8m expressing N2a-P1KO cells with the PIEZO1 activator Yoda2 (*53*) produced discrete Ca^2+^ signals in clusters, which were abolished in the absence of extracellular calcium (Fig. 4G and Supplementary Video 3).

**Fig. 4.**
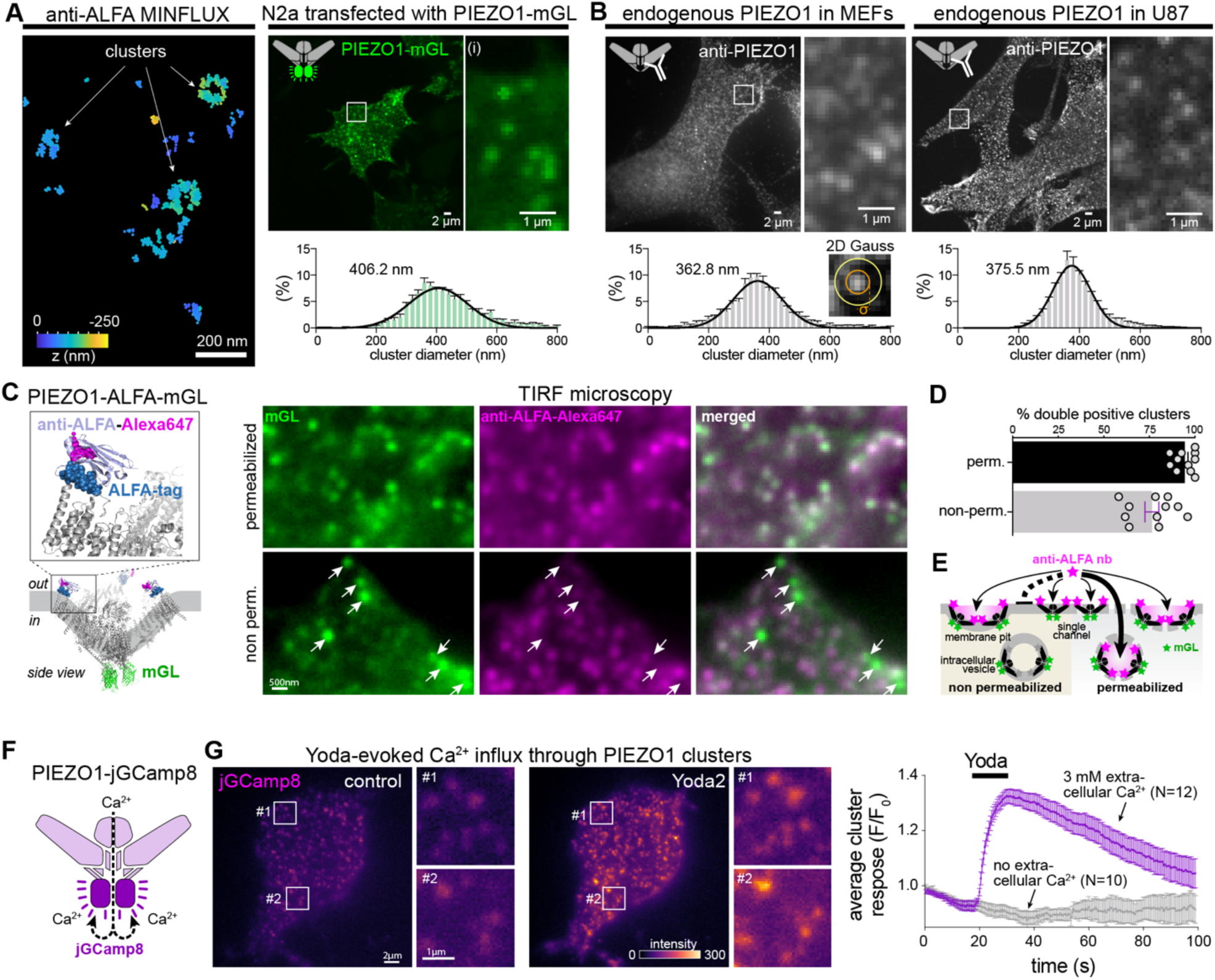
Endogenous and transiently expressed PIEZO1 forms functional cluster at the plasma membrane. **(A)** Example of 3D-MINFLUX raw localization data from extracellular ALFA tag labelled PIEZO1 (left), highlighting the presence of clustered as well as individual channels. Representative TIRF image of a N2a-P1KO cells transfected with PIEZO1-mGL. Insets show closer view of PIEZO1 clusters, which size was determined by a 2D Gauss fit. Distribution of the size of detected clusters are depicted as histogram (bottom) with associated Gaussian fit and average. **(B)** Representative TIRF images (top) of endogenous PIEZO1 clusters labelled with a specific antibody in MEF (left) and U87 (right) cells, with their associated PIEZO1 cluster diameter distribution (bottom). **(C)** Cartoon depicting the position of the extracellular ALFA-tag with its nanobody conjugated to Alexa647 and intracellular mGL in PIEZO1-ALFA-mGL (left). Representative TIRF images of PIEZO1-ALFA-mGL transfected N2a-P1KO cells permeabilized (top) or not permeabilized (bottom), and labelled with Alexa647-coupled nanobody against ALFAtag (right). Scale bar = 500 nm. **(D)** Comparison of the proportion of PIEZO1 clusters double positive for mGL and ALFA-Alexa647 per cell analyzed (N = 11 cells for both conditions, presented as mean ± S.E.M). Two-tailed unpaired t-test, *\*\*\* P* = 0.0003. **(E)** Schematic representation of the accessibility of ALFA-tagged PIEZO1 channels and clusters depending on the permeabilization status of the cell. **(F)** Cartoon depicting the PIEZO1-jGCaMP8m fusion protein. **(G)** Representative TIRF images (left) of N2a-P1KO cells expressing PIEZO1-jGCaMP8m with close-up views on PIEZO clusters before and during Yoda2 application. Time-course of the cell-average cluster normalized fluorescence (F/F0) upon application of Yoda2 with (384 clusters, 12 cells) and without extracellular calcium (174 clusters, 10 cells).

Hence, our data together with previous reports show that endogenously (*26*, *29*, *31–35*) and heterologously expressed PIEZO1 (*20*, *27*, *30*, *33*, *52*) forms clusters that contain functional channels, which possibly contribute to cellular mechanosensitivity (*34*, *38*) (Fig. 4G).

### PIEZO1 clusters form pit-shaped invaginations in the plasma membrane

The unequivocal identification of trimers in densely packed clusters is challenging, because adjacent and partially labelled trimers may produce signal triplets that are indistinguishable from ‘truly’ triple labelled PIEZO1 trimers (Fig. 5A). To circumvent this problem and to obtain more reliable information about the nanoarchitecture of PIEZO1 clusters as well as the absolute number of channels and their relative position within clusters, we next performed 3D-MINFLUX/DNA-PAINT scans using a single-domain nanobody that recognizes the intracellular mGL-tags, which are much closer together such that individual channels are clearly discernible even when partially labelled (Fig. 5A, Fig. S7A-B and Fig. S7C for negative controls). The anti-mGL 3D-MINFLUX localization data perfectly matched the mGL fluorescence pattern in confocal images and resolved individual fluorophores with a localization precision of less than 3 nm in all three dimensions (Fig. 5B-D), which revealed that clusters contain multiple channels and that many channels appear to reside in isolation in the interjacent space (Fig. 5C). To estimate the proportion of isolated channels, we quantified the number of channels that did not have any neighbours within their membrane footprint using previously proposed hypothetical footprint radii between 50 and 100 nm (*6*, *11*, *12*, *54*). This analysis suggested that, depending on the assumed footprint radius, 16.9–44.7 % of the channels are isolated and 32.8–74.5 % only have a single neighbour within their footprint (Fig. 5E), demonstrating that despite the prominence of clusters, a considerable fraction of PIEZO1 channels reside in isolation.

**Fig. 5.**
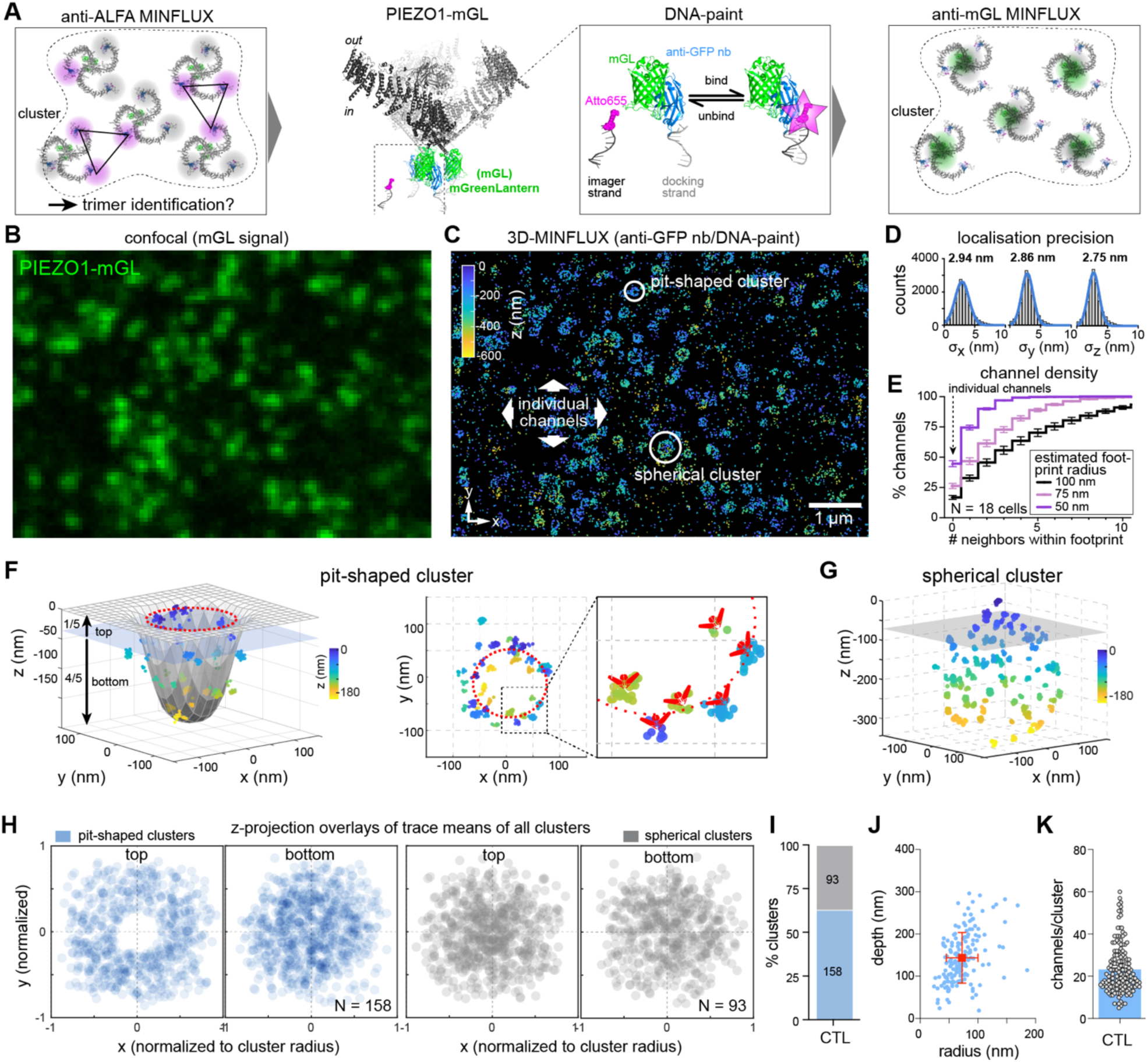
3D-MINFLUX reveals pit-shaped nanoarchitecture of PIEZO1 clusters. **(A)** Cartoon depicting the ALFA-labelled trimers within a PIEZO 1cluster (left). Cryo-EM structure of PIEZO1-ALFA-mGL and localization of C-ter mGreenLantern(mGL)-tag (middle) and the GFP labelling DNA-PAINT approach to improve individual PIEZO1 channel identification within cluster (right). **(B)** Confocal scan of an N2a-P1KO cell expressing PIEZO1-mGL. **(C)** Corresponding 3D-MINFLUX localization data, colored by z depth. **(D)** Distribution of the standard deviations of the MINFLUX traces along the x, y, and z-axis. **(E)** Channel density estimation with the cumulative frequency distribution of the number of PIEZO1 channels neighbors within various footprint radius. **(F)** 3D view (left) and top view (left) of the raw MINFLUX localization data of the pit-shaped cluster cluster marked in (C). The inset shows the putative localization of individual PIEZO1 channels within the cluster. **(G)** 3D view of the raw MINFLUX localization data of the spherical cluster marked in (C). For 360° rotation movies of the clusters shown, see Supplementary Videos S4 and 5. **(H)** Overlay of the 2D projections of the traces means of the top fifth and bottom four fifth (left) of all pit-shaped clusters (blue circles, N=158 cluster from 19 cells) and spherical clusters (grey circles, N=93 clusters from 19 cells). Note, in pit-shaped clusters no channels are present at the center of the top fifth. Bar graph (right) showing the proportions of pit-shaped and spherical clusters.

Most notably, our 3D-MINFLUX scans revealed previously unrecognized details about the nanoarchitecture of PIEZO1 clusters, which appeared as ring-like structures in the 2D-projected MINFLUX data (Fig. 5C). Closer inspection of the MINFLUX localization data in 3D, showed that all clusters exhibited significant expansion along the z-axis, with most clusters having a clearly recognizable pit-like shape, while others appeared to be spherical objects (Fig. 5F, G and Supplementary Videos 4 and 5). To quantify the proportions of pit-shaped and spherical clusters, we segmented the raw MINFLUX localization data of each cluster into signals originating from the upper fifth (top) and the lower four fifth (bottom) (Fig. 5F), and classified clusters in which no channels were present near the center of the top segment as pit-shaped clusters, whereas those clusters that did contain channels in this region were considered as spherical clusters (Fig. 5G). Consistent with the results from the anti-ALFA labelling of unpermeabilized cells (Fig. 4D), this classification scheme suggested that approximately two thirds of the PIEZO1 clusters were pit-shaped and had direct access to the extracellular side (62.9%, 158 from 251), while the remaining third appeared to be spherical and possibly located intracellularly (93 from 251 clusters, Fig. 5H and I). The pit-shaped clusters had a depth of 144 ± 60.1 nm, a radius of 72.8 ± 27.3 nm and contained an average of 23.1 ± 11.3 channels (Fig. 5J and KF).

Previous studies have shown that PIEZO1 enriches in concave environments such as the dimple region of red blood cells (*36*), T-tubules in cardiomyocytes (*32*) and in nanoscale invaginations formed by cells grown on nanobars (*55*), which raises the question if cluster formation results from the accumulation of channels in pre-existing invaginations or if PIEZO1 actively contributes to membrane bending and pit formation. The PIEZO1-pits observed here, were reminiscent of other well-described pit-shaped invaginations, such as clathrin-coated pits, caveolae and intermediate stages of vesicle formation (e.g. COP1 coated cargo vesicles) (*56–58*). To examine if PIEZO1 clusters co-localize with these invaginations, we co-transfected N2a-P1KO cells and MEFs with PIEZO1-mScarlet (Fig. S8) and Clathrin-mGL, and immunolabelled the cells with antibodies directed against Caveolin-1 and COP1, respectively (Fig. 6A and B). The great majority of PIEZO1 clusters did not co-localize with these invagination markers, neither in N2a-P1KO cells nor in MEFs (Fig. 6C). The only exception was a 17.6 % overlap of PIEZO1 clusters with clathrin-coated pits in N2a cells. However, consistent with a prior report (*27*), blocking clathrin-dependent endocytosis with 75µM Dynasore, did not change the density of PIEZO1 clusters in the plasma membrane observed in living cells via TIRF microscopy, suggesting that Clathrin is not required for PIEZO1-pit formation (Fig. 6D). Hence, in summary our data demonstrates, that neither clathrin nor caveolin-1 or COP1 are required for PIEZO1-pit formation, but we cannot rule out that other, yet unknown, accessory proteins are involved. Considering that individual PIEZO1 channels were shown to locally deform the membrane into a dome shape (*12*) and given the remarkable structural similarity between PIEZO1, clathrin and COP1 (Fig. 6E), it is, however, tempting to speculate that PIEZO1 channels themselves drive pit formation due to their curved triskelion structure, which appears to be a common feature of coat proteins that are required for pit formation in other biological processes (*59*, *60*).

**Fig. 6.**
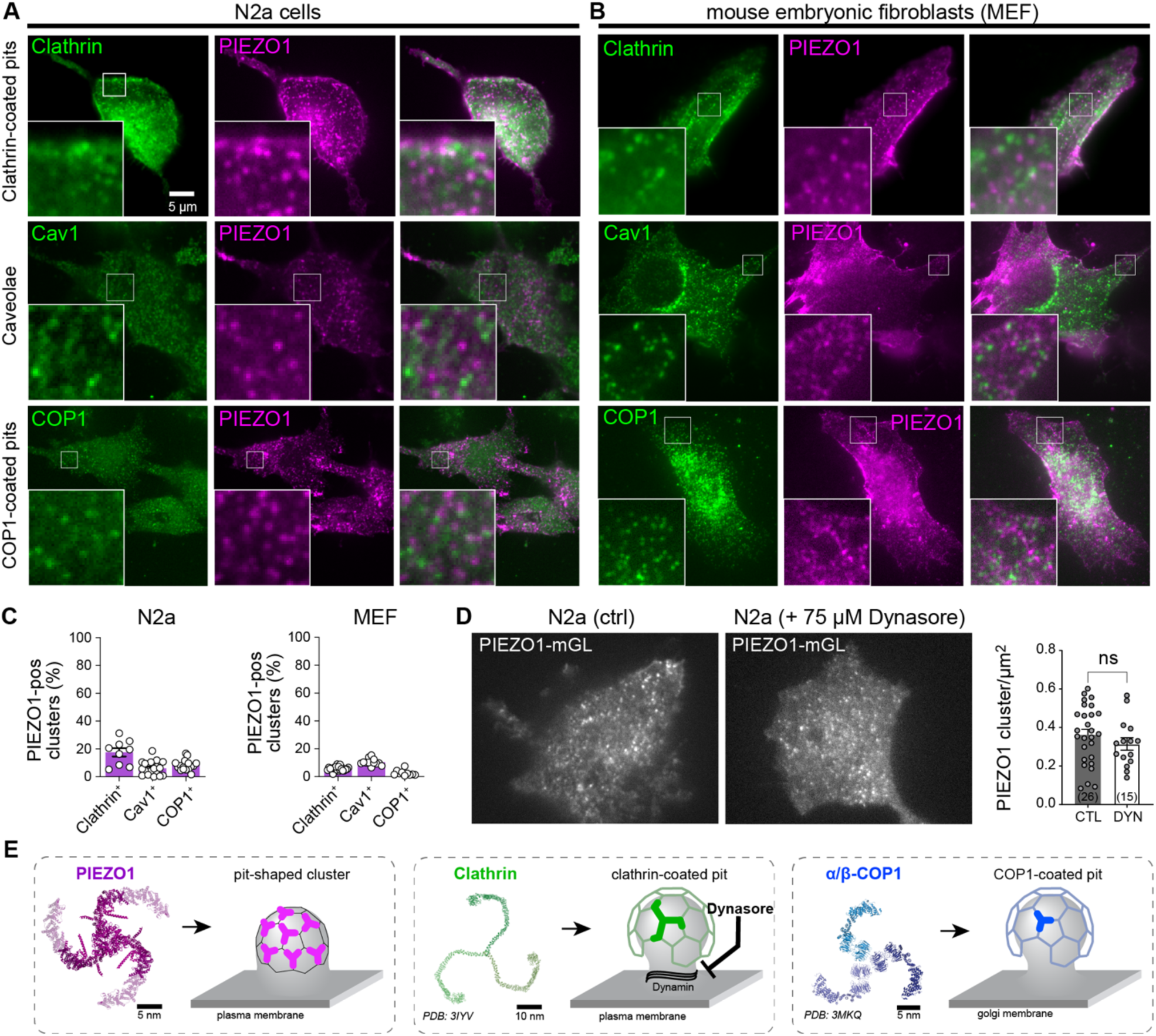
PIEZO1 pit-shaped clusters are distinct cellular invagination structures. **(A-B)**, Representative TIRF images of PIEZO1-mScarlet expressed in N2a-P1KO (**A**) or MEF (**B**) together with Clathrin light chain-mGL to visualize clathrin-coated pits and co-labelled with Cav1 (caveolae) or αCOP1 (COP1-coated pits). **(C)** Quantification of the proportion of PIEZO1 clusters expressing the cellular invagination markers in N2a-P1KO (left) and MEF (right) cells. Data are presented as cell average. **(D)** Representative TIRF images (left) of N2a-P1KO live cells expressing PIEZO1-ALFA-mGL and extracellularly labelled with ALFA-Alexa647 nanobodies with or without the clathrin-dependent endocytosis inhibitor Dynasore. Quantification (right, presented as cell average) of the number of PIEZO1 clusters. Two-tailed unpaired t-test, ns P=0.3141. **(E)** Cartoon depicting the structural similarity of PIEZO1 pit-shaped cluster (left, PDB 7WLT with AFE2JF22F1, shaded) with other pit/invagination-related proteins: Clathrin (middle) and the α/β subunits of the COP1 complex (right).

### Hypo-osmotic stimulation alters PIEZO1 cluster nanoarchitecture

PIEZO1 is activated by changes in membrane curvature and tension evoked by mechanical stimuli such as membrane stretch, cell compression and hypotonic stress-induced cell swelling (*2*, *4*, *5*). We thus examined how the pit-shaped microenvironment of PIEZO1 clusters changes in response to mechanical stimulation, by comparing the PIEZO1 cluster nanoarchitecture of cells fixed during hypo-osmotic stimulation with that of control cells. Exposure of PIEZO1-mGL-expressing N2a-P1KO cells to hypoosmotic stress before and during fixation, significantly reduced the depth of pit-shaped clusters (CTL: 144 ± 60.1 nm vs. OSMO: 110.7 ± 54.8 nm; Fig. 7A-C, Supplementary Videos 6 and 7, Fig. S9), yet the proportions of pit-shaped and spherical clusters were not changed (Fig. 7D). There was also a slight decrease in the average number of channels per cluster, but no change in the pit opening radius (Fig. 7E and F). Considering that PIEZO1 is supposedly activated by changes in membrane curvature and considering further that pit-shaped invaginations exhibit a strong gradient of curvatures ranging from positive values (concave) at the bottom to negative values (convex) at the pit opening (Fig. 7B), small changes in cluster structure or in the relative localization of channels within the pits, could have large effects on the curvature that individual channels are exposed to and hence on their activity. We thus fitted the surface of the clusters and calculated the gaussian curvature of the surface at the coordinates where individual channels were detected. This analysis revealed that the mean curvature that individual channels are exposed to, is significantly reduced during hypoosmotic stimulation and that more channels are exposed to convex curvatures in which they are more likely to be open (Fig. 7G).

**Fig. 7.**
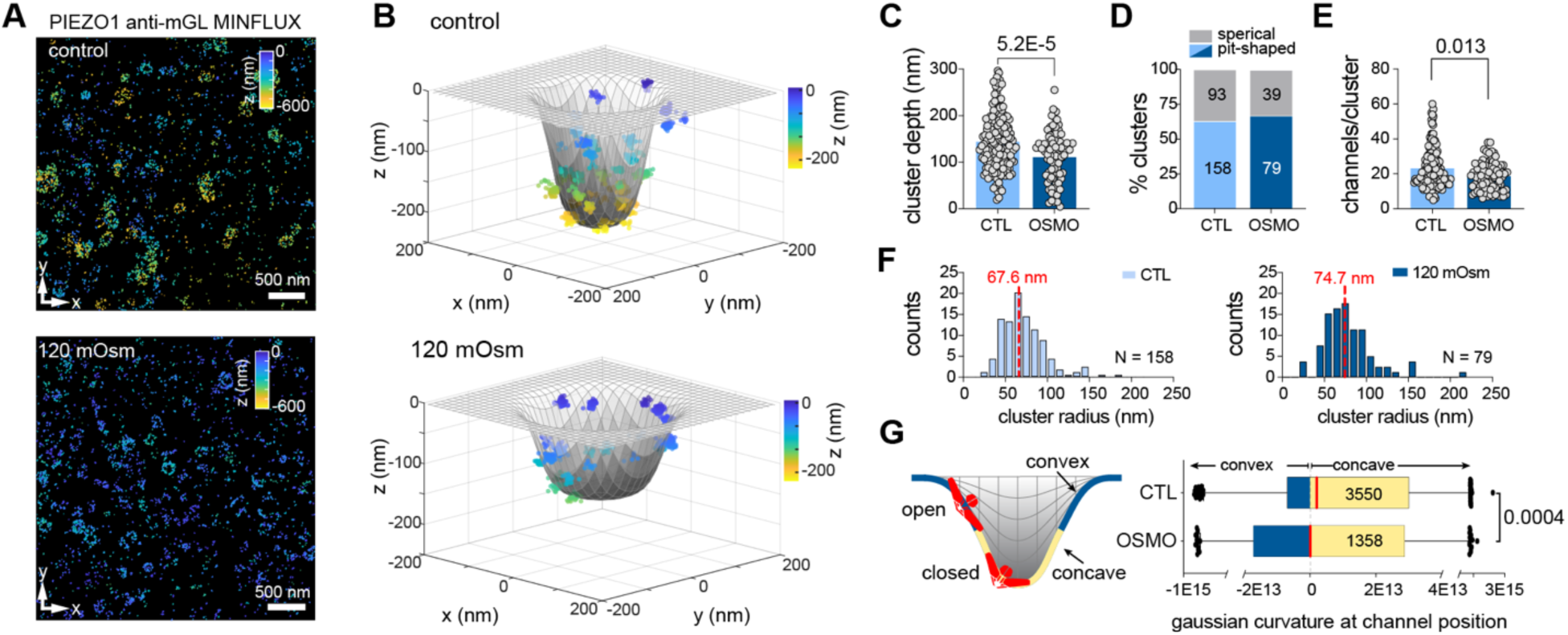
Hypo-osmotic stimulation causes partial flattening of pit-shaped clusters. **(A)** Raw MINFLUX localization of clusters imaged in control cells (top) and challenged with hypoosmotic solution (bottom). **(B)** 3D-views, of the raw MINFLUX localization of representative clusters from control (top) and osmotically stimulated (bottom) cells together with surface fits. **(C)** Comparison of the cluster depths (CTL: N=158 vs. OSMO: N=79, P=0.000052, two-tailed Student’s t-test). **(D)** Proportions of pit-shaped (blue) and spherical clusters (grey) in control (CTL) and hypo-osmotically stimulated (OSMO) cells. Absolute numbers of clusters are within the bar. **(E)** Comparison of the mean ± s.e.m number of PIEZO1 channel found in individual pit-shaped clusters. (CTL: N=158 vs. OSMO:, N=79, P=0.0134, two-tailed Mann-Whitney test). **(F)** Histograms of the radii of all pit-shaped clusters measured at the pit opening. Medians are indicated in red. **(G)** Schematic surface fit in which convex regions are depicted in blue and concave regions are shown in yellow (left). The possible impact of curvature on PIEZO1 (red) conformation is depicted. Gaussian curvatures of the fitted surfaces (see B) calculated at the coordinates where channels are and shown as horizontal box plots, ranging from the lower quartile to the upper quartile. Medians are shown as red lines. Mean curvatures were compared using Mann Whitney test (P = 0.0004, N=3550 for CTL and N=1358 for OSMO).

Together out data suggest that pit-shaped invaginations change their shape in response to hypo-osmotic stimulation resulting in changes in local curvature and possibly activation of individual channels.

## Discussion

A recent MINFLUX study suggested that PIEZO1 is generally more expanded in its native environment than predicted by cryo-EM structures of PIEZO1 reconstituted in liposomes (*23*). Our work extends and refines these findings, by showing that PIEZO1 in intact cells assumes discrete conformations closely matching known cryo-EM structures, and that the preference for specific conformations varies between subcellular compartments (Fig. 1). We also identify the cytoskeleton as a crucial cell intrinsic factor that differentially controls PIEZO1 conformation at rest across subcellular compartments (Fig. 3). Finally, we observed compartment-specific differences in PIEZO1’s susceptibility to Yoda1-induced flattening, which correlate with differences in mechanical and chemical sensitivity (Fig. 2).

Analysis of the interblade distance distribution of PIEZO1 channels in somata using a Gaussian mixture model revealed that interblade distances are not uniformly distributed, suggesting PIEZO1 preferentially adopts discrete and possibly energetically favorable conformations (Fig. 1G). This interpretation is supported by the close match between the interblade distances of the flat and the curved population observed here and those from previously resolved cryo-EM structures (Fig. 1G and H) (*7*). Moreover, a similarly contracted “super-curved” conformation has been described in a recent preprint using single particle cryogenic light microscopy (*61*). Notably, the same three conformations were observed in neurites, but with more channels adopting the super-curved state and fewer appearing flattened (Fig. 1G), suggesting that compartment-specific factors influence PIEZO1 conformation. Membrane curvature, extracellular tethering to the ECM, or channel crowding did not account for these differences (Fig. 3A-C). Disruption of the cytoskeleton with cyto-D, however, altered the distribution of PIEZO1 conformation such that it mirrored the distribution observed in neurites, indicating that differences in cytoskeletal architecture may contribute to the observed compartment-specific differences in PIEZO1 conformation (Fig. 3D-I). This observation is consistent with the idea that PIEZO1 exerts a bending force onto the membrane (*6*, *8*, *11*, *12*), which implicates that PIEZO1 can contract more in cell compartments and nanodomains in which the membrane is more deformable. Considering the membrane and the cytoskeleton as a composite material, the deformability of the membrane at the microscopic level does not only depend on the curvature and lipid composition of the membrane itself but also on cytoskeletal rigidity. Moreover, at the nanoscopic level the membrane is compartmentalized by cytoskeletal attachments, which limit the lateral flow of lipids, such that large membrane compartments created by widely spaced cytoskeletal attachments are more deformable than smaller compartments. Accordingly, modelling studies have proposed that these attachments may impose constraints on the size of PIEZO1s membrane footprint (*6*, *11*) – i.e. the degree of PIEZO1 flattening. Neurites are well known to have a different overall cytoskeletal architecture with less membrane-cytoskeleton attachments (*47–49*) and were shown to be more deformable at the microscopic level (*50*, *62*, *63*). Likewise, cytoskeleton disruption with cyto-D makes the membrane of somata softer (*51*). Moreover, elegant membrane tether pulling studies have shown that the membranes of somata are more resistant to lateral lipid flow than those of axons, possibly due to differences in the density and spatial arrangement of membrane-cortex attachments (*64*, *65*). Hence, together our observations that PIEZO1 is more contracted in neurites and after disruption of the cytoskeleton with cyto-D, support a model in which PIEZO1 conformation at rest is strongly influenced by local membrane deformability, which is controlled by cytoskeletal rigidity and possibly the density of membrane cytoskeleton attachments.

We also assessed how PIEZO1 conformation changes during Yoda1-induced activation. In somata, Yoda1 caused only minor, insignificant interblade expansion (Fig. 2C), as previously reported by Mulhall et al (*23*). In neurites, which are more sensitive to Yoda1 (Fig. 2A) and after cyto-D treatment, which increases the mechanosensitivity of PIEZO1 (Fig. S5), however, Yoda1 induced highly significant and much more pronounced changes in interblade distances (Fig. 2E and 3H). The simple interpretation of these results is that enhanced chemical and mechanical sensitivity correlates with a larger relative change in interblade distance. However, our detailed analysis of the interblade distributions suggested a more complex explanation. Thus, in the somata of control cells, almost 50% of the channels seem to adopt the flat conformation, whereas only 16 % and 30 of the channels were flat in neurites and after cyto-D treatment (Fig. 1G and Fig. 3I). Due to the lack of precise side-chain density in the cryo-EM structure of the flattened conformation, it is unclear if this conformation represents an ‘open’ or an ‘inactivated’ state (*7*, *14*). Considering that 50% of the channels appear to be flat in the soma, yet spontaneous activity is rarely observed in patch-clamp recordings and calcium imaging, our data suggest that most flat channels are in fact in an ‘inactivated’ state. Accordingly, the fraction of channels that are available for activation (i.e. curved and super-curved) is much larger in neurites and after cyto-D treatment as compared to somata, which could explain the observed differences in chemical and mechanical sensitivity. An alternative explanation, at least for the increased Yoda1 sensitivity in neurites, could be that the Yoda1 binding site, which is supposedly located at the intracellular end of the THU8-THU9 interface (Fig. 2B) (*46*), is more accessible when the blades are tilted upward as might be the case in the super-curved conformation, which is more common in neurites as compared to somata (Fig. 1G). Finally, it should be noted that the curved population was almost completely absent in somata challenged with Yoda1 (Fig. 2D), suggesting that most channels transition into the flattened conformation, which possibly represents the inactivated state. In neurites and after cyto-D treatment, by contrast, the proportions of the curved population were not altered by Yoda1, but, notably, their mean interblade distances increased by more than 2 nm (Fig. 2F and 3G), suggesting that many of these channels transitioned into the partially flattened conformation, which has an estimated interblade distance of ∼24.7 nm (*14*). The partially flattened conformation represents an intermediate open state with a significantly dilated pore and thus more channels in this conformation could explain the increase sensitivity of neurites and cyto-D-treated somata.

In summary, our data highlight that PIEZO1 is not a rigid, binary switch but a conformationally plastic channel whose resting and activated conformations are dynamically tuned by the physical properties of its microenvironment, positioning the cytoskeleton as a key modulator of mechanosensitive signaling.

Accordingly, our findings have significant implications for understanding compartmentalized mechanotransduction in neurons and other polarized cells and serve as a mechanistic framework for future studies that should aim to identify the molecular nature and spatial organization of membrane-cytoskeleton attachments that regulate PIEZO1 conformation.

In addition to investigating how the conformation of individual PIEZO1 channels is shaped by the native cellular environment and during channel activation, we also examined the subcellular distribution of PIEZO1 and the impact of clustering on membrane topology. Previous studies using immunolabeling, live-cell calcium imaging, and STORM super-resolution microscopy have reported the presence of PIEZO1 clusters in both endogenous (*26*, *29*, *31–35*) and heterologous systems (*20*, *27*, *30*, *33*, *52*). Using 3D-MINFLUX/DNA-PAINT nanoscopy, we corroborate and expand upon these findings, by showing that the majority of PIEZO1 clusters form pit-shaped invaginations in the plasma membrane (Fig.5C-I). The interpretation that these invaginations are contiguous with the plasma membrane rather than intracellular vesicular structures, is supported by the observation that Yoda1-induced Ca^2+^ signals originating from PIEZO1 clusters require extracellular calcium (Fig. 4G) and by our extracellular ALFA-tag labelling experiments (Fig. 4C-E). Moreover, we found that mechanical stimulation via hypo-osmotic stress caused significant flattening of these pit-like structures and Gaussian curvature mapping revealed a redistribution of PIEZO1 channels within the pits upon hypo-osmotic challenge, such that more channels are exposed to convex curvature (Fig. 7), a condition previously shown to facilitate channel opening (*8*, *66*). Hence, our data suggest that the curvature gradient along the pit axis offers an energetically favourable microenvironment for mechanotransduction, where small changes in pit shape could alter the local mechanical landscape experienced by individual channels.

With regards to the effect of clustering on PIEZO1 structure and function, in-silico modelling studies had suggested that the opposing curvatures of the membrane footprints of two nearby channels would create an energetic constraint in the interjacent membrane such that nearby channels would either repel each other or induce mutual flattening to reduce the overall energy of the system (*6*, *12*, *39*). Our data contradict these models and demonstrate that clustering instead causes the formation of pit-shaped invaginations that accommodate up to twenty channels and more (Fig. 5). PIEZO1 appears to prefer concave membrane environments, as it was shown to accumulate in the dimple region of red blood cells (*36*), cardiomyocyte T-tubules (*32*), and artificially induced membrane curvatures (*55*), which raised the question if the PIEZO1-pits observed here resulted from the accumulation of PIEZO1 in pre-existing invaginations or if PIEZO1 itself actively contributed to their formation. The lack of co-localisation of PIEZO1 with clathrin, Cav1 and COP1 together with the insensitivity of cluster density to pharmacological inhibition of clathrin-mediated endocytosis (Fig. 6), demonstrate that PIEZO1 pits are distinct from classical membrane invaginations such as clathrin-coated pits, caveolae, or COPI-associated vesicles, which supports the latter idea. Moreover, considering that PIEZO1 exerts a strong bending force onto membranes (*7*, *8*, *11*), pit formation appears to be an effective mechanism to minimize the energy of the PIEZO1 cluster system, by allowing the membrane footprints of individual PIEZOs to seamlessly integrate into the overall pit curvature without causing energetic constraints in the membrane between neighbouring channels PIEZOs. Finally, even though we cannot definitively rule out a contribution of other, yet unidentified, membrane bending proteins to pit formation, the remarkable structural resemblance of PIEZO1 with the well-described coat proteins clathrin and COP1 (Fig. 6E), which are known to sculpt membrane pits, strongly supports the intriguing possibility that PIEZO1 itself might actively drive the formation of membrane pits.

Collectively, our findings suggest a model in which PIEZO1 clusters are not passive aggregates but rather functionally and structurally specialized units that create distinct microenvironments within the plasma membrane. These clusters may act as localized mechano-responsive microdomains capable of amplifying mechanical stimuli via structural changes in their topology. Thus, our work provides a framework for future studies that should address the molecular determinants of cluster formation, the role of lipid microdomains in organizing these structures, and whether accessory proteins contribute to the formation and stabilization of pit-shaped PIEZO1 clusters.

## Materials and Methods

### Generation of PIEZO constructs

A mouse PIEZO1-IRES-GFP plasmid (Addgene #80925) was used as the initial template to generate most of the constructs of the present work, using a similar strategy described in earlier studies (*20*, *52*). PIEZO1-mGreenLantern fusion protein (PIEZO1-mGL) was generated by amplifying the coding sequence of the green fluorescent protein mGreenLantern(*67*) from a LifeAct-mGreenLantern plasmid (gift from G. Petsko, Addgene #164459), and inserted with a SG linker after the C-ter of the PIEZO1 plasmid, where the IRES-GFP sequence has been excised beforehand by PCR. PIEZO1-mScarlet was generated with restriction enzymes, by digesting the PIEOZ1-IRES-GFP plasmid with BspEI and FseI enzymes (New England Biolabs) and a PCR-amplified mScarlet fragment from a pAAV-CAG-FLEX-mScarlet plasmid (gift from Rylan Larsen, Addgene #99280). Ligation (T4 ligase, Promega) was performed overnight at 16°C. The PIEZO1-ALFA-IRES-GFP construct was generated by inserting a synthetic DNA fragment coding for amino acid 1 to 180 of mouse PIEZO1, with the ALFA tag sequence inserted after amino acid position H86 and flanked by one proline on each side (GeneArt Custom Gene Synthesis, ThermoFisher)(*68*), into a PIEZO1-IRES-GFP plasmid excised by PCR for the corresponding 1-180 coding region. A plasmid version of PIEZO1-ALFA without the IRES-GFP sequence but with a C-ter HA-tag was also generated, using a previously generated PIEZO1-HA template(*69*). The PIEZO1-ALFA-IRES-GFP plasmid was later used as a template to generate the PIEZO1-ALFA-mGreenLantern (PIEZO1-ALFA-mGL), by exchanging the IRES-GFP sequence with mGreenLantern, as described above. The PIEZO1-jGCaMP8m fusion was generated by inserting at the C-ter of PIEZO1 the coding sequence of jGCaMP8m(*70*) (a gift from GENIE Project, Addgene #162372) with a GSGG linker, following a similar and validated strategy described previously(*38*). To facilitate visualization of transfected cells due to the low basal fluorescence of the jGCaMP8m fusion, a second plasmid was generated by adding a P2A-mScarlet sequence after the jGCaMP8m. All DNA fragments were generated by PCR (primers from Sigma) using KAPA HiFi polymerase (Roche) and cloning was performed with homologous recombination (Gibson Assembly, NEBuilder HiFi, New England Biolabs). PCR reactions were digested with DpnI (New England Biolabs, 37°C, 1h) and column purified with standard kits (NucleoSpin from Macherey-Nagel or PureLink from Invitrogen) before being assembled by homologous recombination and then transformed in electrocompetent Stbl4 or Dh5a bacteria (Invitrogen) and grown at 30°C for 48h (Stbl4) or 37°C overnight (Dh5a). Selected clones were entirely sequenced (Eurofins) to ensure that no mutation was present.

### Cell culture

Mouse neuroblastoma Neuro-2a PIEZO1-Knockout cell line (N2A-P1KO) was generated and characterized previously from Neuro-2a ATCC CCL-131 (a gift from G.R Lewin(*43*)). Cells were grown in Dulbecco’s Modified Eagle Medium (DMEM) and optimal Minimal Essential Medium (opti-MEM) (1:1 mixture), supplemented with 10% Fetal Bovine Serum, 2mM L-glutamine and 1% peniciline/streptomycine (all from ThermoFisher). U87 (U87-MG, ATCC HTB-14) and Mouse Embryo Fibroblast (MEF) (ATCC SCRC-1008) were grown in DMEM, supplemented with 10% Fetal Bovine Serum and 1% peniciline/streptomycine. For some live TIRF imaging experiments, medium without phenol red was used. Cells were cultured at 37°C with 5% CO2. Cells were seeded on poly-L-lysine (PLL, Sigma) coated and methanol- and acid-washed glass coverslips (12mm diameter for patch-clamp recordings, #1.5 and 18mm diameter for Minflux and TIRF imaging on fixed samples), or PLL-coated 35mm glass-bottom dishes (MatTek High Precision #1.5 Coverslip, TIRF microscopy live-imaging). Cells were transfected one or two days after plating using polyethylenimine (PEI, Linear PEI 25K, Polysciences). For one 12mm coverslip, 7μl of a 360μg/ml PEI solution is mixed with 9μl PBS. Plasmid DNA is diluted in 20µl PBS (0.6 μg/coverslip) and then the 16μl PEI-PBS solution is added to the DNA solution. After at least 10 minutes of incubation at room temperature, the DNA-PEI mix is added drop by drop and mixed by gentle swirling. For a 35mm dish or 18mm coverslip, 2.0μg DNA is used and PBS/PEI volumes are adjusted accordingly. 24h later, the medium is replaced by fresh one. In selected experiments investigating neurites, N2A-P1KO cells were serum-starved for approximately 12h to promote neurite outgrowth. Cells were then used within 24h to 48h.

### Electrophysiology

All PIEZO constructs generated were tested for proper functionality in patch-clamp assays and compared to control PIEZO1-IRES-GFP. Mechanically activated currents were recorded at room temperature using EPC10 amplifier with Patchmaster software (HEKA Elektronik). Borosilicate patch pipettes (2–6 MΩ for whole-cell, 1.5–3.5 MΩ after fire-polishing for cell-attach) were pulled with a P-97 Flaming-Brown puller (Sutter Instrument). For whole-cell patch clamp, intracellular buffer contained the following: 125mM K-gluconate, 7mM KCl, 1mM MgCl_2_, 1mM CaCl_2_, 4mM EGTA, 10mM HEPES (pH 7.3 with KOH). For single-channel cell-attach: 130mM NaCl, 5mM KCl, 1mM MgCl_2_, 1mM CaCl_2_, 10mM HEPES, 10mM TEA-Cl (pH7.3 with NaOH). The control bath solution for whole-cell contained the following: 140mM NaCl, 4mM KCl, 1mM MgCl_2_, 2mM CaCl_2_, 4mM glucose and 10mM HEPES (pH 7.4 with NaOH). For single channel recordings, the bath solution contained: 140mM KCl, 1mM MgCl_2_, 2 CaCl_2_, 10 Glucose, 10 HEPES (pH7.4 with KOH). Cells were held at a holding potential of −60 mV (whole-cell and cell-attach). Mechanical stimulation in whole-cell experiments was done with a series of 15 mechanical stimuli in 0.6 μm increments with a fire-polished glass pipette (tip diameter 2-3μm) that was positioned opposite to the recording pipette, at an angle of approximately 45° to the surface of the dish and moved with a velocity of 1 μm/ms by a piezo driven micromanipulator (Preloaded Piezo actuator P-840.20, Physik Instrumente). Negative pressure stimuli in cell-attach experiments were applied for 500ms with the High-Speed Pressure Clamp device (HSPC; ALA Scientific Instruments), with −5mmHg increments up to −80mmHg. A pre-pulse of +5mmHg was applied before negative-pressure stimuli to improve recovery from inactivation (*66*). For I/V experiments, pressure stimulus was adjusted on a cell-by-cell basis to optimally evoke single-channel openings.

The evoked whole cell currents were recorded with a sampling frequency of 200 kHz and filtered with 2.9 kHz low-pass filter. Pipette and membrane capacitance were compensated using the auto function of Patchmaster. Recordings with excessive leak currents, unstable access resistance and cells which giga-seals did not withstand at least 7 consecutive mechanical steps stimulation were excluded from analyses. Mechanical thresholds of PIEZO currents were determined by measuring the mechanical stimulus that evoked the first sizeable peak current, defined as the point in which the current significantly differed from the baseline (more than 6 times the standard deviation of the baseline). The inactivation time constants (τ_inact_) were measured by fitting the mechanically activated currents with a single exponential function (C1+C2*e ^(–(t–t0)/τinact)^, where C1 and C2 are constants, *t* is time and τ_inact_ is the inactivation time constant. For each cell, only peak currents between 100pA and 1500pA were used for τ_inact_ calculation and averaged from cell to cell.

The evoked cell-attach currents were recorded with a sampling frequency of 50 kHz and filter with a 2.9 kHz low-pass filter. Maximal pressure-evoked currents over the course of a given stimulus (I) were normalized to the absolute maximal response of the cell at any pressure (I_max_). Normalized pressure-response curve (I/I_max_) from individual cells were fitted with a Boltzmann sigmoid to determine individual P50 (in mmHg).

Single-channel amplitudes at a given holding potential (−140mV to −40mV, 20mV steps) were determined as the difference between the peaks of the gaussian fits of the trace histogram over multiple 1s segments. Unitary conductance was determined from the linear regression fits of the I/V plot of individual cells. Recordings with excessive leak currents or unstable baseline were excluded. Recordings that displayed non-inactivating responses or unstable openings were also not used for further I/V analyses. All electrophysiology analysis was performed in IgorPro (Wavemetrics) using custom scripts.

### Calcium imaging of neurites

N2A-P1KO cells were plated on 12mm coverslips, transfected with PIEZO1-mScarlet (0.6 μg per coverslip) together with a CMV-jGCaMP8m vector (0.2 μg per coverslip) and serum starved to promote neurite outgrowth. Cells were washed once with PBS and incubated with a calcium imaging buffer containing 140mM NaCl, 4mM KCl, 1mM MgCl_2_, 3mM CaCl_2_, 4mM glucose and 10mM HEPES (pH 7.4 with NaOH). Fluorescent images were acquired every second (500ms exposure time) on an Olympus BX40 upright microscope equipped with standard Quad filter (Chroma), fluorescent lamp (HBO 100) and shutter (Lambda 10-2, Sutter Instrument) with a 40x water-immersion objective (LUMPlanFl/IR, Olympus), visualized with a CoolSnap HQ2 camera (Photometrics) and acquired with the MetaFluor software (Molecular Devices). Perfusion and fast solution exchanged was achieved with a gravity-driven perfusion system (ValveLink8.2, AutoMate Scientific). Cells were first perfused with control solution for at least 20 seconds before being exposed to Yoda1 (Sigma, 100nM, 300nM or 1µM) for 20 seconds, followed with a washout with control solution. Only one field of view per coverslip was used to avoid Yoda1 leakage. Only cells that are double positive for PIEZO1 and jGCaMP8m and that have neurites were considered for analysis. Cell body and associated neurites were segmented in ImageJ and the time course of normalized fluorescence ratio (F/F_0_) was calculated as the ratio between the jGCaMP8m fluorescence intensity (a.u) at a given time (F) and the average fluorescence intensity per PIEZO-mScarlet transfected cell averaged over a 10s interval during the initial control perfusion (F_0_). Within the Yoda1 perfusion time, the maximal F/F_0_ ratio per cell was extracted. For a cell having multiple neurites, values were averaged and compared to its cell body with paired test.

### Preparation of samples for TIRF imaging

For live samples, N2A-P1KO cells were plated on glass bottom dishes as described before and transfected with PIEZO1-ALFA-mGL (live labelling) or PIEZO1-jGCaMP8m (calcium imaging of PIEZO clusters). Two to three days after transfection, samples were processed for live labelling and imaging. For live labelling, cells were washed once with phenol-red free medium and incubated for 3min at 37°C with a nanobody against ALFA tag conjugated to Alexa-647 (FluoTag-X2 anti-ALFA, NanoTag Biotechnologies, RRID: AB_3075981) diluted 1:100 in cell culture medium. Cells were then washed twice and imaged immediately. Labelled live samples were kept for no more than 20 minutes. To evaluate the impact of clathrin mediated endocytosis on the number of PIEZO clusters in living cell, the dynamin inhibitor Dynasore (Sigma, final concentration 75µM) was pre-incubated for 2 hours before and kept during the image acquisition. For calcium imaging, cells were incubated in a buffer containing 140mM NaCl, 4mM KCl, 1mM MgCl_2_, 3mM CaCl_2_, 4mM glucose and 10mM HEPES (pH 7.4 with NaOH) and then perfused for 20 seconds with a 2.5µM solution of Yoda2 (Sigma). In experiments with no extracellular calcium, the 3mM CaCl_2_ was replaced by 5mM EDTA.

For fixed samples, N2A-P1KO, U87 or MEF cells were plated on 18mm coverslips. In experiments investigating accessibility of PIEZO cluster, N2A-P1KO cells were transfected with PIEZO1-ALFA-mGL. In experiments investigating PIEZO cluster and their co-localization with different markers, N2A-P1KO, U87 and MEF cells were transfected with PIEZO1mScarlet or PIEZO1mScarlet together with a plasmid encoding Clathrin Light Chain fused to mGL (Clc-mGL, gift from G. Petsko, Addgene #164462) to visualize clathrin coated pits. In experiments investigating native PIEZO1 clusters in MEF and U87, cells were not transfected. Two to three days after transfection or plating (native PIEZO1 clusters), cells were washed once with PBS and fixed for 10min at room temperature with a mixture of 1% PFA and 0.1% glutaraldehyde in PBS. Fixative was removed and quenched with 5mM NaBH_4_ in PBS (one quick wash and another one for 5min at room temperature) and then with 50mM glycine and 50mM NH_4_Cl in PBS (one quick wash and another one for 5min at room temperature). Samples were further washed thrice with PBS for 5min at room temperature. In experiments evaluating accessibility of PIEZO clusters in N2A-P1KO, cells were then blocked with either a mix of 5% FBS, 1% BSA in PBS (No permeabilization) or with additionally 0.2% Triton 100X and 0.05% Tween 20 (Permeabilization) for 30min at room temperature. Samples were then labelled with the ALFA nanobody conjugated to Alexa-647 diluted 1:100 into respective blocking buffers and incubated for 1 hour at room temperature. For co-labelling experiments and exploration of native PIEZO1 clusters, cells were processed with the permeabilized buffer and then incubated overnight at 4°C with either a rabbit antibody against Cav1 (1:200) (abcam, RRID:A B_303405), a rabbit antibody against COPIα (1:1000) (ThermoFisher 23GB3485, RRID: AB_3249045) or a rabbit antibody against PIEZO1 (1:100) (Novus NBP1-78446, RRID: AB_11020328) diluted in permeabilized buffer. Samples were then washed 3 times for 5min at room temperature with their respective blocking buffers. A post-fixation step was performed for ALFA nanobodies, using the same fixative mixture as before, for 5min. Fixatives were quenched and then washed 3 more times with PBS. For co-labelling or native PIEZO1 labelling, samples were incubated for 1 hour at room temperature with a donkey anti-rabbit antibody coupled to Alexa647 (1:1000) (Life Technologies). After several washes, coverslips were then mounted on slides with ProLong Glass Antifade Mountant (ThermoFisher). Samples were then imaged within 2 days.

### TIRF microscopy imaging and acquisition

TIRF imaging was performed on a Nikon Eclipse TiE microscope and with a Roper iLAS2 TIRF module. The objective was an oil immersion Nikon CFI Plan Apo Lambda 100x (NA 1.45) and the camera used was a Photometrics Prime 95B back-illuminated sCMOS, having a resolution of 1200 x 1200 and with a pixel size of 11 µm, giving a final pixel size of 0.110µm. A 1.5x magnification lens was added for some experiments, giving a final pixel size of 0.073µm. Cells were illuminated with solid-state lasers of 488nm, 560nm and 640nm (20% power). Acquisition was done with VisiView software (version 5.0.0, Visitron Systems). For live imaging, an incubation chamber (okolab) was used to adjust temperature (37°C), CO2-concentration (5%) and humidity. Live cells were imaged for 30s with a frame rate of 10Hz (approximately 100ms exposure time per frame) for live labelling experiments, or for 2min with a frame rate of 2Hz (calcium imaging).

### TIRF microscopy analysis

Due to the mobility of PIEZO clusters in live cell, clusters were tracked for analysis as described before(*20*, *52*). For calcium imaging experiments, time-lapse recordings were first preprocessed in ImageJ with a bleach correction (simple ratio(*71*)). PIEZO track analysis was then performed with TrackMate v7.13.2(*72*, *73*)and the following parameters: DoG detector, blob diameter of 0.35µm, spots quality filter value of 0.10-1, simple LAP tracker with a linking distance of 0.7µm, a gap closing distance of 0.7µm and a maximal gap closing frame number of 2. The average intensity over time within each detected cluster was then extracted and normalized to the baseline, defined as the average intensity over the last 10 seconds of acquisition before Yoda2 was applied. For live labelling experiments, clusters were detected and counted in the first frame with TrackMate using the same parameters as above.

For fixed samples and experiments investigating the extracellular accessibility of the cluster, analysis was performed in ImageJ. PIEZO clusters were identified on the mGL signal (488nm excitation): a background subtraction (rolling ball radius 5) was first performed, a Gaussian blur was applied (sigma 2 and 3) and blurred images were subtracted. An Auto-local Threshold was then used (Bernsen method, radius 5) followed by a Particle Detection (min. size = 10 and max. size =50 pixels) to identify clusters as Region of Interests (ROIs). Intensity of the mGL and ALFA-Alexa647 signals were then measured within the ROIs and mGL clusters were classified or not as ALFA-Alexa647 positive. A similar approach was used to investigate colocalization of PIEZO1mScarlet cluster with the selected cellular markers. For fixed samples and experiments investigating endogenous PIEZO1 clusters and their size (Figure 4), clusters were detected with TrackMate as ROIs and their coordinates were extracted, imported in IgorPro and fitted with a 2D Gaussian fit to estimate their diameter.

### Preparation of samples for DNA-PAINT Minflux imaging

N2A-P1KO cells were plated on 18mm coverslips as described in previous section and transfected with PIEZO1-mGL, PIEZO1-ALFA-HA or PIEZO1-ALFA-mGL. Two to three days after transfection, cells were processed for fixation as described above for TIRF imaging or first serum starved, treated with hypotonic solution, Yoda1 or cytochalasin D and then fixed as described above. For hypo-osmotic experiments, cells were washed once with PBS, then incubated for 3min at 37°C in a 120mOsm Ringer’s solution (48.8mM NaCl, 5mM KCl, 10mM HEPES, 10mM glucose, pH 7.4) and finally fixed for at least 15min at room temperature in a hypotonic fixative (0.5% PFA, 0.05% glutaraldehyde, 200mOsm). For cytochalasin D (Sigma) experiments, cells were incubated for 20min at 37°C with 2µM cytochalasin D diluted in culture medium, which was kept after in the fixative solution (Minflux imaging) or in the extracellular buffer (electrophysiology). For Yoda1 experiments, cells were incubated for 3min at 37°C with 50µM Yoda1 diluted in culture medium, which was kept after in the fixative solution. After quenching and washing of fixatives, samples were blocked with Antibody Incubation buffer (Massive Photonics) for 30min at room temperature. The cells were incubated for 1h at room temperature or overnight at 4°C with Massive-Tag-Q Anti-ALFA or Anti-GFP nanobodies conjugated with a DNA docking strand (both from Massive Photonics) diluted at 1:100 (Anti-ALFA) or 1:200 (Anti-GFP) in Antibody Incubation buffer. PIEZO1-ALFA-HA samples were fixed, blocked and permeabilised as described above for TIRF samples, then incubated with the anti-GFP nanobody and with a rabbit anti-HA antibody (1:500, ThermoFisher, RRID: AB_2533988). Cells were washed thrice with 1X Washing buffer (Massive Photonics) and then post-fixed for 5min. Fixatives were quenched and washed as described before. PIEZO1-ALFA-HA samples were then incubated for 1 hour at room temperature with a secondary antibody donkey anti-rabbit coupled to Alexa488 (1:1000) (Life technologies), and further washed with PBS. Samples were then incubated for 10 min with 100-200 µl of gold nanoparticles for future stabilization (gold colloid 250nm, BBI Solutions, or A12-40-980-CTAB, Nanopartz). Unbound nanoparticles were rinsed extensively with PBS and remaining nanoparticles were further stabilized with PLL for at least 1 hour at room temperature. Cells were then washed thrice with PBS before mounting. Labelled samples were used within 3 days. DNA-PAINT imagers (Massive Photonics) conjugated to Atto 655 were freshly diluted in Imaging buffer (Massive Photonics) for a final concentration of 1 ∼ 2nM (Atto 655 and ALFA imaging, Imager sequence #3) or 1nM (Atto655 and GFP imaging, Imager sequence #2). A drop of imager dilution was added into a cavity slide and coverslips were mounted and sealed with picodent twinsil (picodent).

### 3D Minflux imaging

Minflux imaging was performed on an Abberior MINFLUX commercial microscope built on an inverted IX83 microscope with a 100X UPlanXApo objective (Olympus) and using Imspector Software (Abberior Instruments). Daily alignment and calibration of the excitation beam pattern and position of the pinhole was performed using fluorescent nanoparticles (abberior Nanoparticles, Gold 150 nm, 2C Fluor 120 nm, Abberior Instruments). Cells were identified with a 488nm confocal scan and the transient binding of imagers with Atto655 was quickly verified with 640 nm confocal scan. At least 2 gold fiducials were present in the field of view and used by the active-feedback stabilization system of the microscope (IR 975nm laser, Cobolt, and CCD camera, The Imaging Source), having typically a precision below 1nm in all three axes and being stabled for hours. A ROI of 1×1 to 5×5 µm (up to 8×8 µm for some overnight recordings) was selected at the cell-coverslip interface, except for some neurites where focus was set approximately in their middle height. Laser power in the first iteration and pinhole were set at 16% laser power and 0.83 A.U pinhole. Final laser power in the last iteration is scaled up by a factor of six. ROIs were imaged for at least 2 hours and up to overnight (∼12 hours) using the standard 3D Minflux sequence (Supplementary Table 1). Detection for Atto 655 signals was performed with two avalanche photodiodes channels (650-685nm and 685-720nm) that were pooled. Specificity of the nanobodies and imagers used in this study and of the Minflux signal was tested by incubating samples transfected with PIEZO1-mGL with the anti-ALFA nanobody and samples transfected with PIEZO1-ALFA-HA with the anti GFP nanobody (Fig. S2A-C and S7C).

### Minflux data analysis

Final valid localizations from Minflux iterations were exported from Imspector software as matlab files. Custom Matlab scripts were then used for post-processing and filtering of the data, as well as subsequent operations and data visualization. Data filtering involved first an efo filter (effective frequency at offset, kHz, retrieved for each individual valid locations) based on its overall distribution for each individual measurements: a threshold was then selected to filter-out potential multiple emitters. An additional cfr filter (center frequency ratio) was used for mGL/GFP cluster experiments, with a cutoff value of 0.5 (0.8 elsewhere, directly implemented during the acquisition). Then, localizations from the same emission trace, i.e. with the same trace identification number (TID), having a standard deviation of more than 10 nm in the x, y, z axes and less than 3 (ALFA) or 5 (GFP) localizations were excluded (see Fig. S2D). For ALFA signals, filtered traces were trimmed of their first 2 localizations, as they are often apart from the rest and the majority of the localization cloud, possibly caused by diffusing imager molecules. For GFP signals, localizations for each trace were aggregated (group of 3), as described elsewhere(*74*). Remaining traces were further corrected for the refractive index mismatch between the coverslip and the sample, applying a scaling factor of 0.7 for all traces in the z dimension(*40*).

For mGL/GFP cluster experiments (Fig. 5 and 7), clusters were semi-manually selected based on a first round DBSCAN (variable parameters adjusted with imaging signal density, typically epsilon of 80 to 160nm and minPoints of 100 raw localizations) and analysed as follows. The center of mass of each individual filtered trace (i.e trace with the same TID) within the clusters was calculated. Due to the nature of DNA-PAINT labelling and the proximity of individual mGL molecules at PIEZO C-ter, neighbouring signals coming from the same PIEZO channel were estimated and averaged using a DBSCAN on the trace average calculated before, with a minPoints of 2 and epsilon of 25nm (based on recording precision and labelling error), ultimately giving a position of an individual PIEZO channel (see Fig. S7A and B). To classify clusters into pit-shaped (open at the top) and spherical clusters, we segmented the raw MINFLUX localization data of each cluster into signals originating from the upper fifth (top) and the lower four fifth (bottom) (Fig. 5F), and classified clusters in which no channels were present near the center of the top segment (i.e. within a distance of 0.25 times the radius from the centroid), as pit-shaped clusters, whereas those clusters that did contain channels in this region were considered as spherical clusters (Fig. 5G). To estimate the local curvature of the membrane at the position of individual PIEZO1 channels within cluster, we fitted the 3D distribution of the trace means using a modified 2D Gaussian distribution that contained a term correcting for cluster width, Z = top + depth –depth/(1+e^((2*width^2 – ((X – CenterX)^2) – ((Y – CenterY)^2))/slope^2)), where X, Y, Z are the coordinates of the channels, CenterX and CenterY are the arithmetic means of the X and Y coordinates of all channels. For surface fitting ‘top’ (Z-coordinate of the topmost channel) and ‘depth’ (distance between the topmost and the lowest channel) were held constant and the width and slope of the clusters were fitted using the custom fit function in IgorPro. Fitting was performed using the Levenberg-Marquardt least orthogonal distance method implemented in IgorPro8 (wavemetrics) that is based on the ODRPACK95 code (*75*). The gaussian curvatures at the coordinates of the channels were then calculated using a custom written script in Matlab (see code availability).

For ALFA trimer experiments, signals were filtered and the center of mass of each individual trace was calculated. Traces originating from repeated detection of the same protomer were identified using DBSCAN clustering with minPoints of 2 and epsilon of 8 nm. The 8 nm cutoff was chosen based on the localisation precision of MINFLUX and the possible ALFA-tag flexibility (Fig. 1B). The position of the protomers that were detected multiple times was determined by calculating the mean coordinate of the localisations clustered by the DBSCAN algorithm. PIEZO1 trimers in which all three protomers were labelled and detected by MINFLUX were identified by searching for three adjacent traces that were less than 40 nm apart, which we assumed is the maximum distance two Atto-655 molecules bound to the same PIEZO1 trimer can possibly have based on available flattened PIEZO1 structures, and that had no other neighbouring traces within a distance of 60 nm (Fig. S3. Moreover, only trimers in which the maximum interblade angle was smaller than 120° were considered. Interblade distance for a given trimer was calculated as the average of the three protomers distance. Interblade distance distribution was fitted with a gaussian mixture model in Matlab. Further visualization of trimers was done by a 2D in-plane projection of the raw localizations for each protomer, followed by a fit with a bivariate Gaussian distribution and displayed with its probability density.

### Structure modelling and data visualization

For visualization purposes, full length PIEZO1 in different putative conformations were generated. The predicted AlphaFold structure of mouse PIEZO1 (AF-E2JF22-F1-v4) was aligned and superposed onto the experimentally determined curved (PDB 6B3R), flat (PDB 7WLU) and intermediate-flat (PDB 8IXO) PIEZO1 structure to visualize the unresolved peripheral blade. The final constructs display the experimentally resolved structure with the added missing blade parts from AlphaFold as transparent color (Fig. 1H). A full length PIEZO1 trimer bearing an ALFA tag at position H86 and a C-ter GFP together with their respective nanobody (PDB 6I2G for ALFA, PDB 3K1K for mGL/GFP), DNA docking site, imager and fluorophore was also generated. All subsequent modification operations and visualization were performed in PyMol (version 2.5.5, Schrodinger). Plasmid design and visualization was performed with SnapGene (version 7, Dotmatics). All the other data, graphics and schematics were elaborated and visualized in Matlab, IgorPro, Illustrator (Adobe) and GraphPad Prism (version 10, GraphPad Software).

### Statistical tests and reproducibility

All experiments in this study were performed independently at least three times, yielding similar results. No statistical method was used to predetermine sample size. Experiments were not randomized and investigators were not blinded during experiments and analysis. Data distribution was systematically evaluated using D’Agostino–Pearson test and parametric or non-parametric tests were chosen accordingly. The statistical tests that were used, the exact P-values and information about the number of replicates are provided in the display items or the corresponding figure legends.

## Acknowledgements

We thank Dr. Antonio Failla and the team of the imaging core facility at UKE Hamburg (DFG Research Infrastructure Portal #RI_00489) for technical assistance with the MINFLUX microscope. Funding for the MINFLUX microscope was awared by the the Hamburgische Investitions- und Förderbank (IFB, grant no. 51164232) under the Operational Programme Hamburg ERDF 2014-2020, REACT-EU of the European Regional Development Fund (ERDF). We also thank Mr. Haider Al-Marsoomi and Ms. Claudia Lüchau for assistance with cloning and cell culture.

## Author contributions

C.V. performed all imaging and patch clamp experiments, wrote code and analyzed the data. L.R. performed patch-clamp experiments and wrote analysis code. N.Z. performed patch-clamp experiments. S.G.L. conceptualized the study, acquired funding, wrote analysis codes, analyzed data and wrote the manuscript.

## Data and materials availability

All data supporting the article, such as the MINFLUX and patch-clamp analysis outputs for each experimental condition, are provided as source data. Raw data and reagents that are not commercially available are available from the corresponding author upon reasonable request. Source data are provided with this paper. The custom Matlab codes for Minflux analysis are available at https://github.com/clementverkest/PIEZO1_MinfluxAnalysis.

## Supplementary Materials

Supplementary Figure S1-S9

Supplementary Table S1

Supplementary Videos S1-S7 sample

