## SupplementaryMaterial for "Cluster nanoarchitecture and structural diversity of PIEZO1 at rest and during activation in intact cells"

Verkest *et al.*

**This PDF file includes:**

Figs. S1 to S9
Tables S1

**Other Supplementary Materials for this manuscript include the following:**

Movies S1 to S7

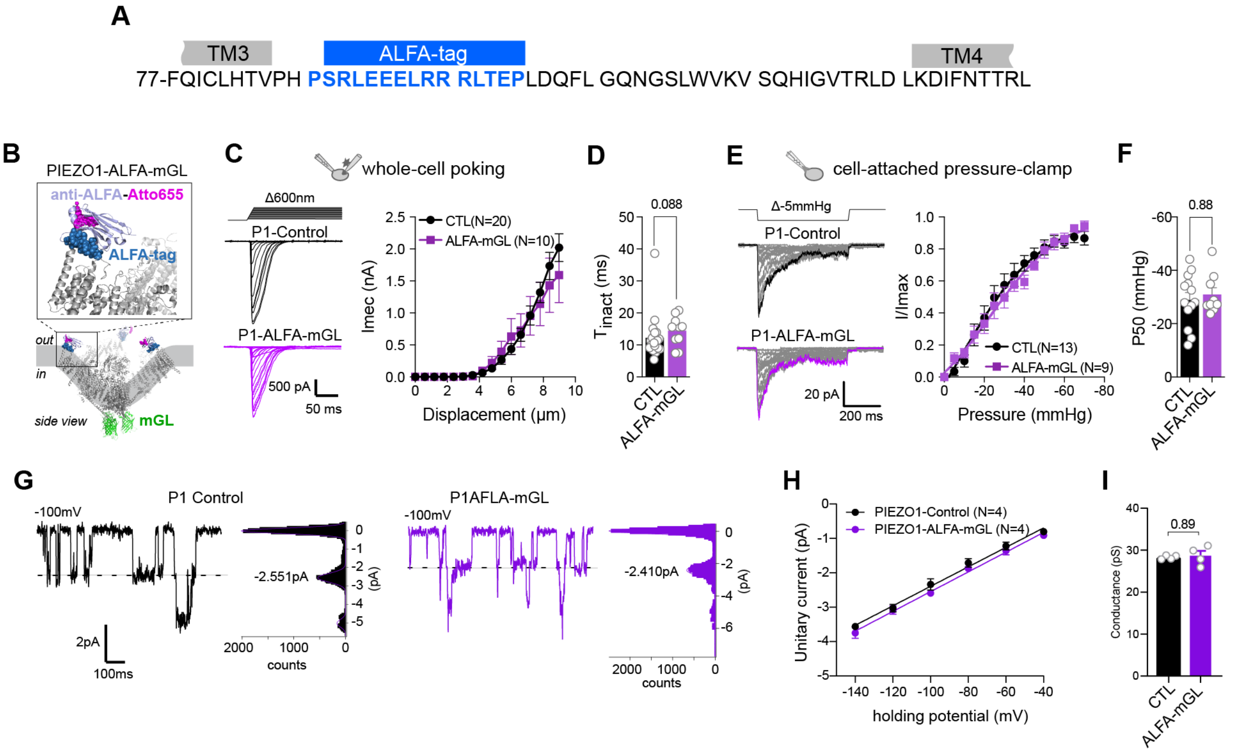


**Fig. S1 | Functional characterization of PIEZO1-ALFA-mGreenLantern.** **(A)** Amino acid sequence of PIEZO1 indicating the position of the inserted ALFA-tag. **(B)** Cartoon depicting the position of the extracellular ALFA-tag and intracellular mGL in PIEZO1-ALFA-mGL. **(C)** Example traces (left) of poking-evoked whole-cell currents of the indicated channel variants (PIEZO1-IRES-GFP, Control, top and PIEZO1-ALFA-mGreenLantern, bottom) in N2a-P1KO and displacement-response curve of mean ± s.e.m. peak current amplitudes (right). **(D)** Comparison of the mean ± s.e.m of the inactivation time constants of PIEZO1 currents using two-tailed Mann-Whitney. **(E)** Representative traces (left) of stretch-evoked cell-attached currents and normalized pressure-response curve of mean ± s.e.m. peak current amplitudes ($I/I_{max}$) of PIEZO1-Control and PIEZO1-ALFA-mGL. **(F)** Comparison of the mean ± s.e.m. P50 values with two-tailed unpaired t-test. **(G)** Example traces of stretch-evoked single channel openings (left) and corresponding all-points histogram (right), with Gaussian fits and indicated peak average of PIEZO1-IRES-GFP (Control) and PIEZO1-ALFA-mGL. **(H)** Relation between mean ± s.e.m. single-channel amplitudes and holding potential fitted with a linear regression. **(I)** Comparison of the mean ± s.e.m. single-channel conductance with Mann Whitney test.

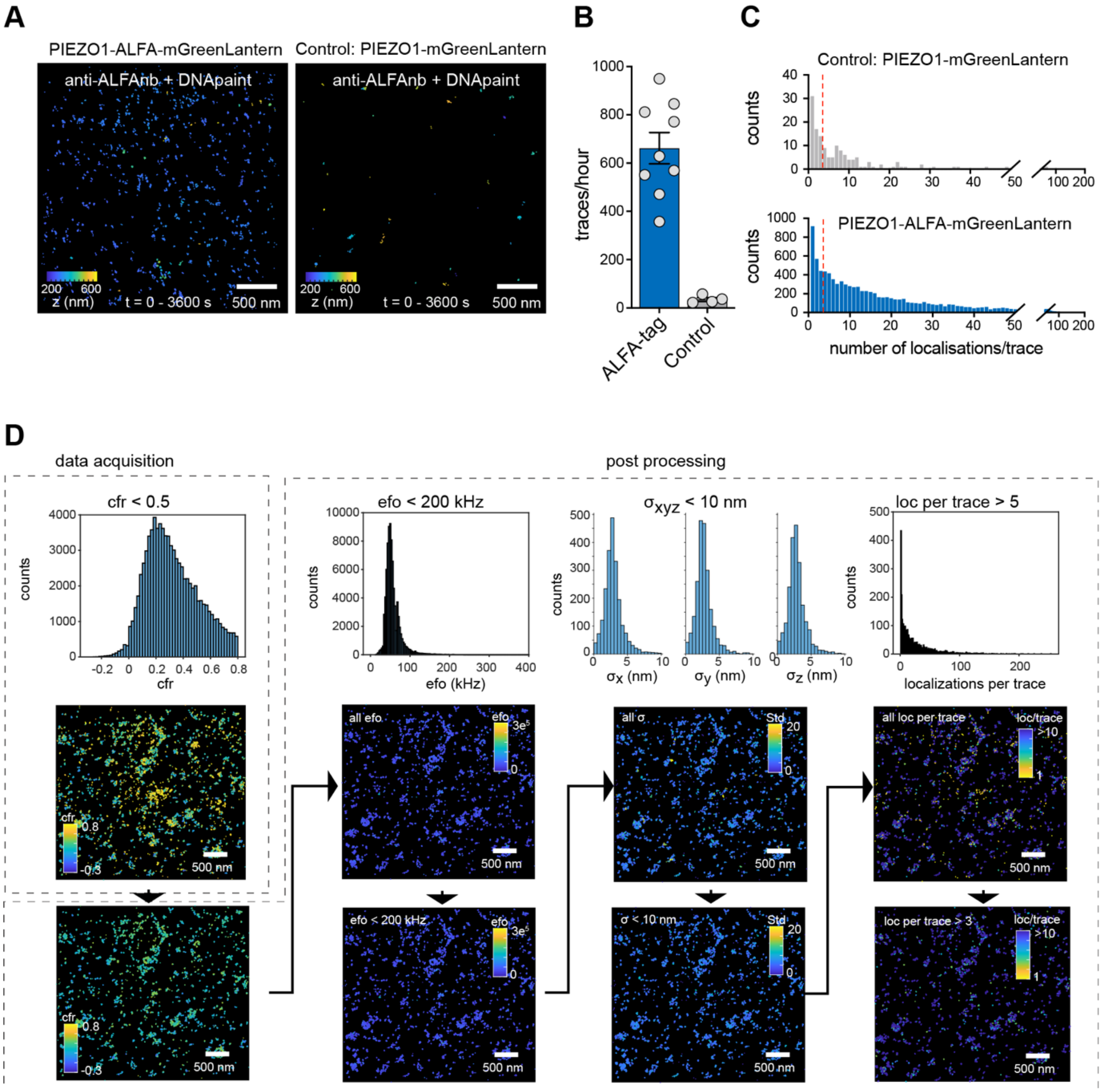


**Fig. S2 | 3D MINFLUX DNA-PAINT acquisition and processing pipeline of PIEZO1-ALFA-mGreenLantern data. (A)** Example of raw MINFLUX DNA-PAINT localizations from a PIEZO1-ALFA-mGL transfected N2a-P1KO cell labelled with an anti-ALFA nanobody and imaged for 2 hours with corresponding imager conjugated to Atto655 (left) and from a PIEZO1-mGL transfected cell labelled with anti-ALFA (right) and imaged in the same condition. Localizations are colored according to their z-position. **(B)** Bar graph of the average number of MINFLUX traces per hour detected for the two conditions: PIEZO1-ALFA-mGL with anti-ALFA, N=9 cells versus PIEZO1-mGL with anti-ALFA, N=4 cells. Comparison with Mann-Whitney test, $P$=0.0028. **(C)** Histogram of the number of localizations per detected trace for Control (PIEZO1-mGL with anti-ALFA, N=4, top) and PIEZO1-ALFA-mGL (N=9, bottom). The red dashed line indicates the cutoff value of 3 used for filtering of the data. **(D)** Overview of the 3D-MINFLUX data filtering pipeline.

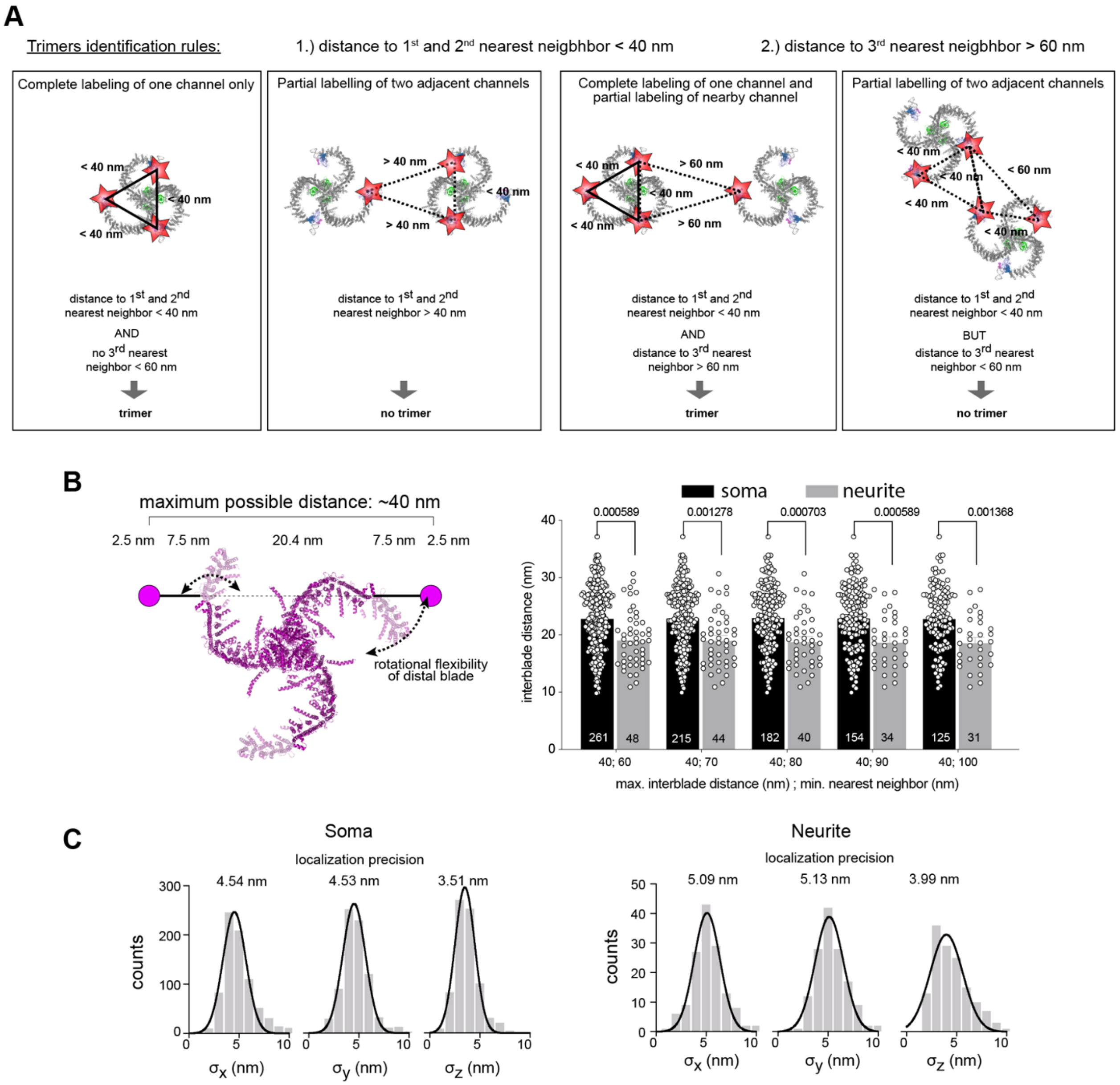


**Fig. S3 | PIEZO1-ALFA-mGreenLantern trimer identification. (A)** Overview of the PIEZO1 trimers identifications rules from MINFLUX ALFA signals. **(B)** Schematic representation of a full length flattened PIEZO1 structure (left), illustrating the maximum measurable interblade distance when taking into account peripheral blade flexibility and linkage error. Bar graphs (right) of the mean interblade distance between soma- and neurite-localized PIEZO1 trimers detected with different cutoffs for trimer identification: maximum interblade distance of 40nm and varying minimum nearest neighbour distance. Comparison with Mann Whitney-tests. **(C)** Distribution of the standard deviations along the x, y, and z-axis of the MINFLUX traces making soma (left) or neurite (right) identified PIEZO1 trimers.

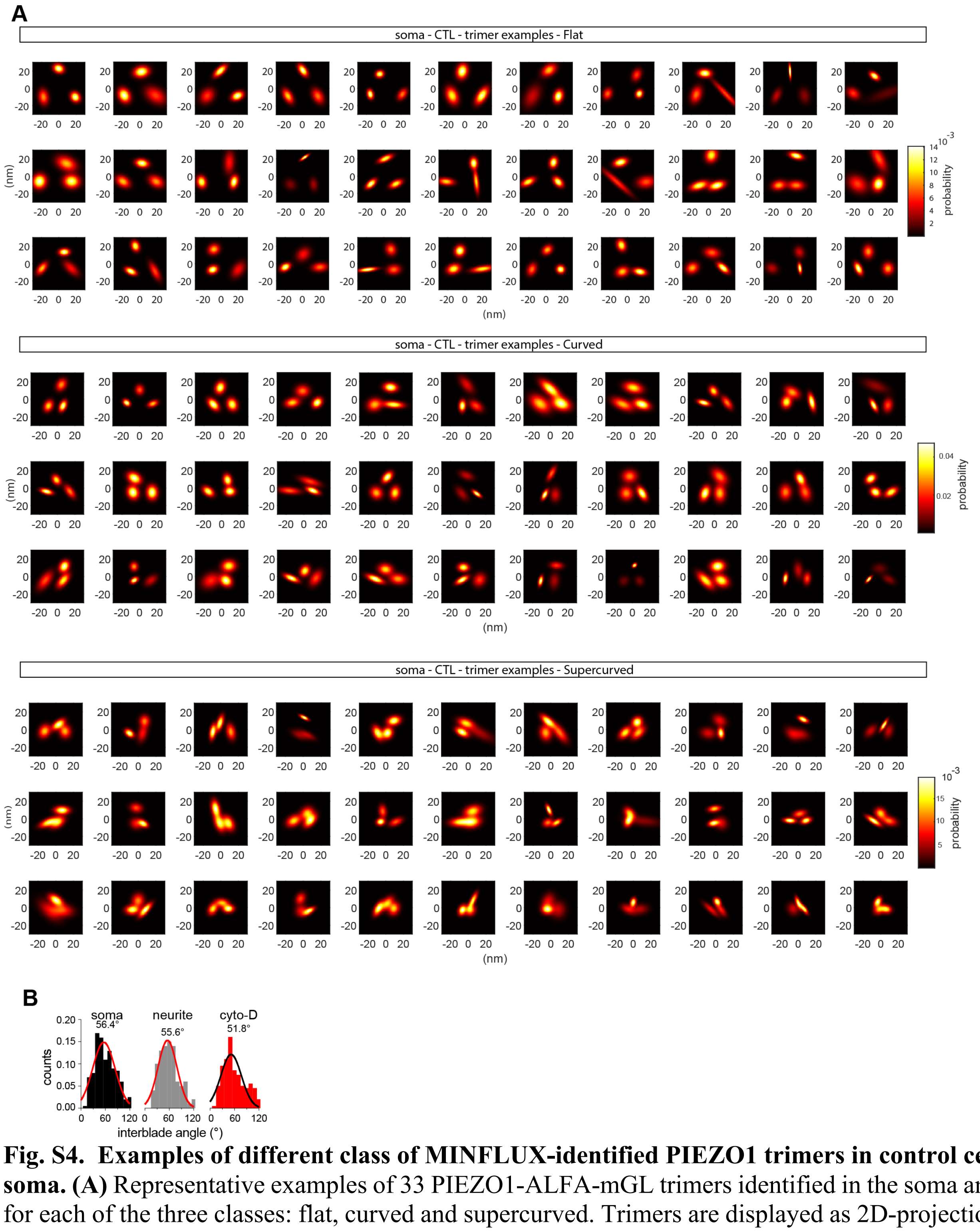


**Fig. S4. Examples of different class of MINFLUX-identified PIEZO1 trimers in control cell soma. (A)** Representative examples of 33 PIEZO1-ALFA-mGL trimers identified in the soma and for each of the three classes: flat, curved and supercurved. Trimers are displayed as 2D-projection of the probability density of the raw localizations making each protomer. **(B)** Histogram of all the measured interblade angle for soma- (black), neurite- (gray) and cytoD-identified PIEZO1 trimers (red). Gaussian fit (red or black line) and average are indicated on the graph.

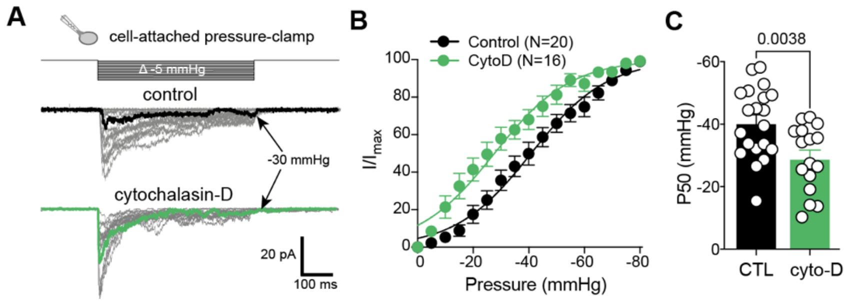


**Fig. S5 | Effect of cytoskeletal disruption by cyto-D on PIEZO1 cell-attached currents. (A)** Example traces of PIEZO1-mediated currents evoked by negative pressure in cell-attached patch-clamp recordings with or without cyto-D. **(B)** Normalized (I/Imax) pressure-response curves of PIEZO1 currents recorded from control (black circles) and cyto-D treated (green circles) cells. **(C)** Comparison of the mean ± sem P50 values, using Student's t-test (CTL: p50 = -40.6 ± 2.5 mmHg, N=20 vs. cyto-D: p50 = -29.1 ± 2.6 mmHg, N =16; P = 0.0038). White circles show values from individual cells.

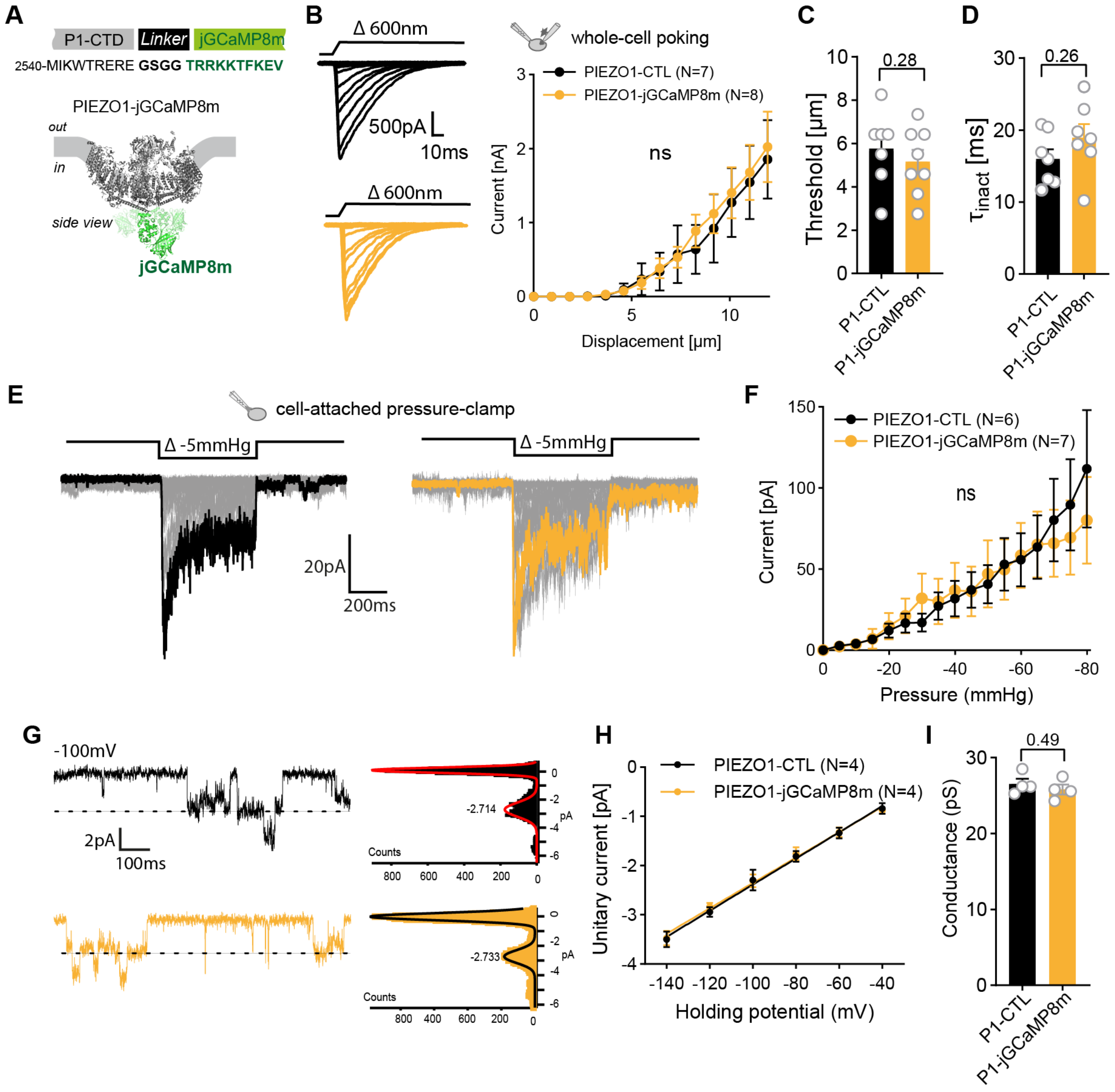


**Fig. S6 | Functional characterization of PIEZO1-jGCaMP8m. (A)** Amino-acid sequence of the inserted jGCaMP8m (top) and its putative location at PIEZO1 C-ter structure (PDB 7st4). **(B)** Example traces (left) of poking-evoked whole-cell currents of the indicated channel variants (PIEZO1-IRES-GFP, top and PIEZO1-jGCaMP8m, bottom). Displacement-response curve (right) of mean peak current amplitudes. **(C)** Comparison of the mean mechanical thresholds using Mann-Whitney test. **(D)** Comparison of the mean of the inactivation time constants of PIEZO1 currents using Mann-Whitney test. **(E)** Representative traces of stretch-evoked cell-attached currents from PIEZO1-Control (left) and PIEZO1-jGCaMP8m (right). **(F)** Pressure-response curve of mean peak current amplitudes of PIEZO1-Control and PIEZO1-jGCaMP8m. **(G)** Example traces of stretch-evoked single channel openings (left) and corresponding all-points histogram (right), with Gaussian fits. **(H)** Relation between mean single-channel amplitudes and holding potential, fitted with a linear regression. **(I)** Comparison of the mean single-channel conductance with Mann-Whitney test.

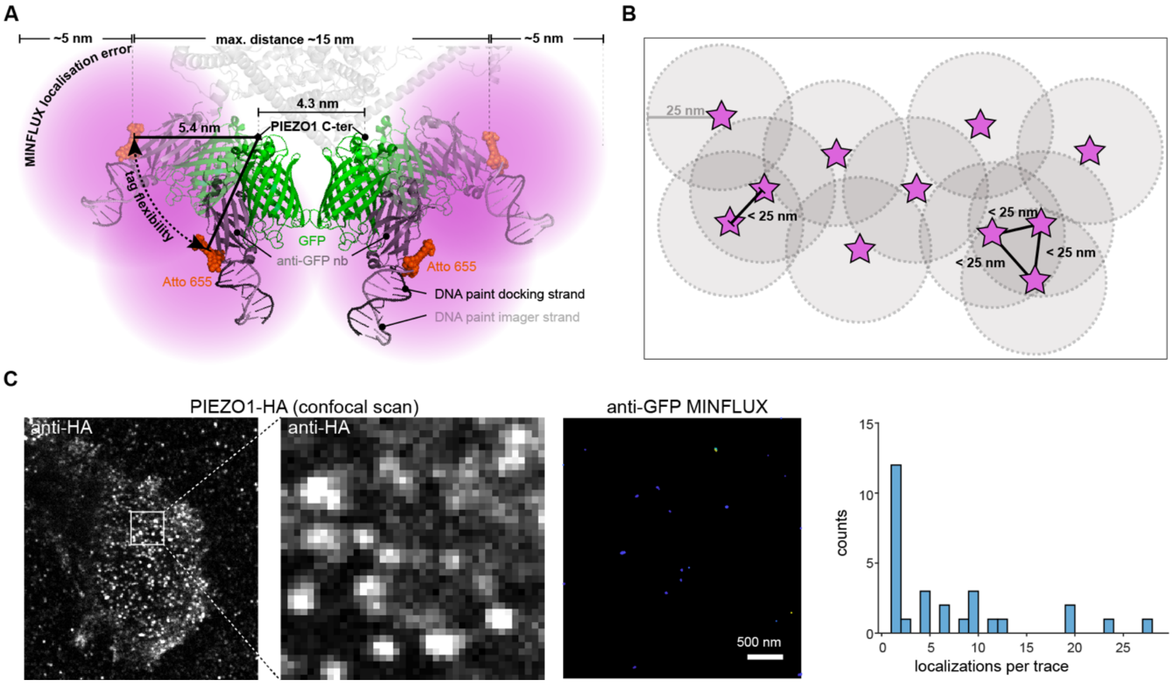


**Fig. S7 | 3D MINFLUX acquisition and processing pipeline of PIEZO1mGreenLantern data.** (**A**) Cartoon showing the extend and the sources of linkage error of PIEZO1-mGL/GFP MINFLUX recording. **(B)** Assignment rules for GFP MINFLUX signals to individual PIEZO channel. **(C)** Selectivity of the GFP nanobody and MINFLUX signals. Representative confocal scan of PIEZO1-ALFA-HA in N2a-P1KO cell (left), labelled with the anti-GFP nanobody used for MINFLUX DNA-PAINT together with an anti-HA antibody for cluster visualization, with a closeup of the selected region for MINFLUX scan (middle, 3x3µm). Samples were imaged with 1nM of GFP-nb corresponding imager conjugated to Atto655. Raw MINFLUX localizations are depicted (right), together with the corresponding histogram of the number of localizations per trace.

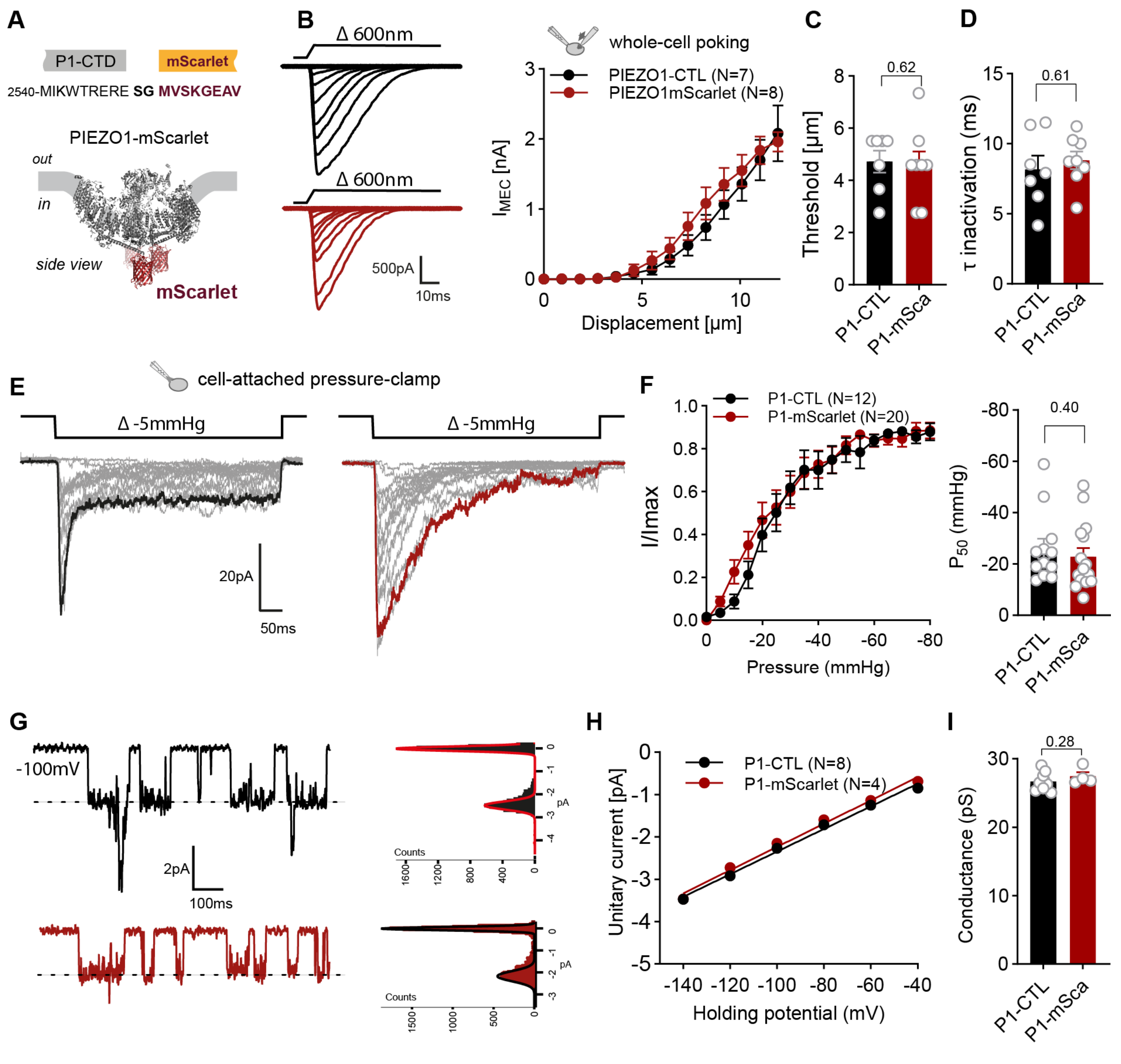


**Fig. S8 | Functional characterization of PIEZO1-mScarlet. (A)** Amino-acid sequence of the inserted mScarlet (top) and its putative location at PIEZO1 C-ter structure (PDB 5lk4). **(B)** Example traces (left) of poking-evoked whole-cell currents of the indicated channel variants (PIEZO1-IRES-GFP, top and PIEZO1-mScarlet, bottom). Displacement-response curve (right) of mean peak current amplitudes. **(C)** Comparison of the mean mechanical thresholds using Mann-Whitney test. **(D)** Comparison of the mean of the inactivation time constants of PIEZO1 currents using Mann-Whitney test. **(E)** Representative traces of stretch-evoked cell-attached currents from PIEZO1-Control (left) and PIEZO1-mScarlet (right). **(F)** Pressure-response curve of normalized mean peak current amplitudes ($I/I_{max}$) of PIEZO1-Control and PIEZO1-mScarlet (left). Bar graph of mean P50 (right). Comparison with Mann-Whitney test. **(G)** Example traces of stretch-evoked single channel openings (left) and corresponding all-points histogram (right), with Gaussian fits. **(H)** Relation between mean single-channel amplitudes and holding potential, fitted with a linear regression. **(I)** Comparison of the mean single-channel conductance with Mann-Whitney test.

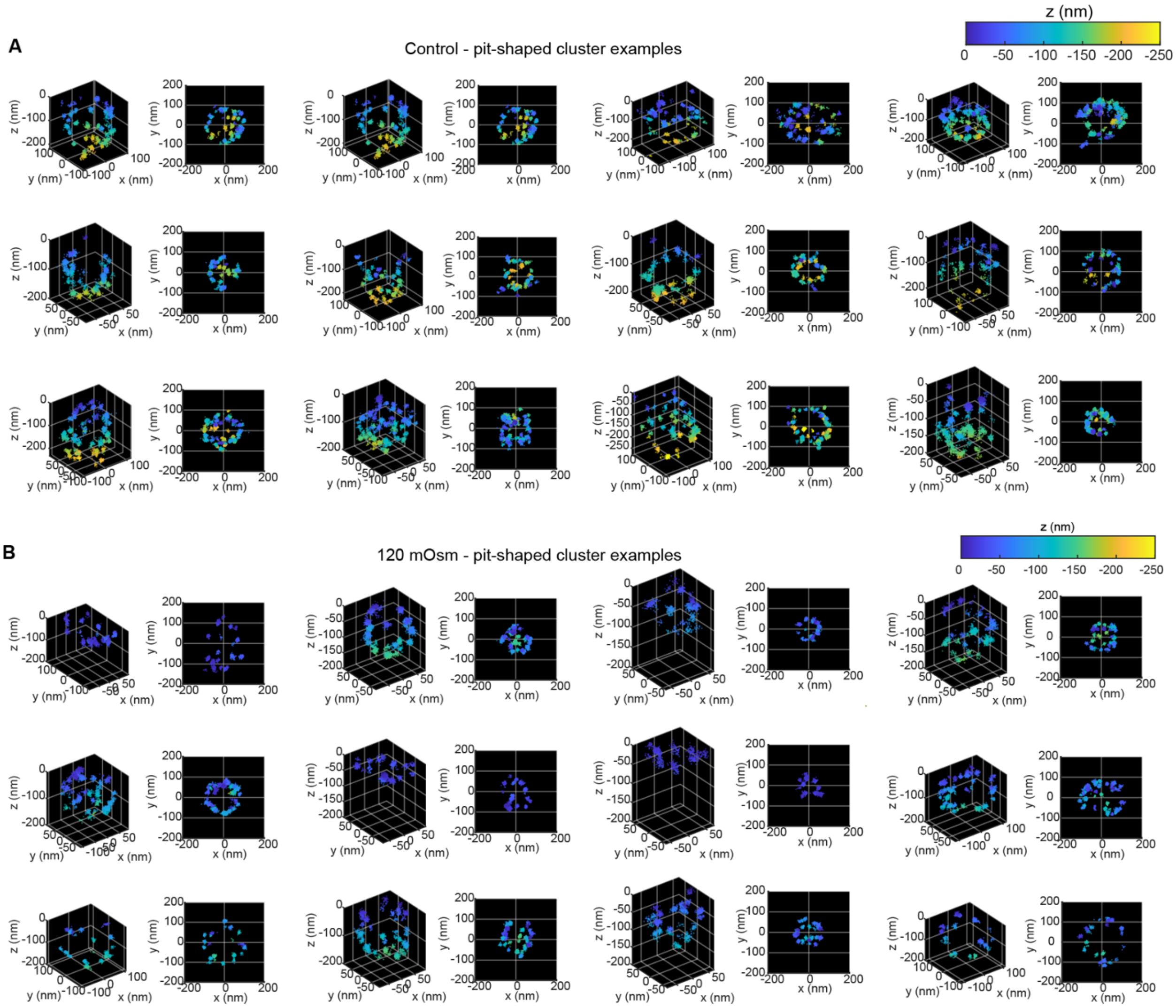


**Fig. S9 | Examples of control and osmotically-challenged pit-shaped PIEZO1 cluster. (A)** Representative examples of 12 control PIEZO1 pit-shaped clusters. MINFLUX raw localizations making the cluster are depicted in a 3D view (left), alongside a z-projected view (right). Localizations are colored based on the relative z position to the top of the cluster. **(B)** Representative examples of 12 pit-shaped osmotically activated PIEZO1 clusters.

| Iteration | Modality | Pattern diameter (nm) | photon limit | back-ground limit | dwell time (ms) | pattern repeat | Stickiness | CFR limit | laser power factor |
|---|---|---|---|---|---|---|---|---|---|
| 0 | hexagonal | 251 | 160 | 15000 | 1 | 1 | - | none | 1 |
| 1 | zline | 251 | 400 | 15000 | 1 | 1 | 2 | none | 1 |
| 2 | square | 251 | 100 | 10000 | 1 | 5 | 2 | none | 1 |
| 3 | zline2 | 251 | 50 | 10000 | 1 | 5 | 2 | none | 1 |
| 4 | square | 132 | 67 | 10000 | 1 | 5 | 2 | 0.9 | 2 |
| 5 | zline2 | 132 | 33 | 10000 | 1 | 5 | 2 | none | 2 |
| 6 | square | 66 | 67 | 10000 | 1 | 5 | 2 | 0.8 | 4 |
| 7 | zline2 | 66 | 33 | 10000 | 1 | 5 | 2 | none | 4 |
| 8 | square | 35 | 100 | 10000 | 1 | 5 | 2 | none | 6 |
| 9 | zline2 | 35 | 50 | 10000 | 1 | 5 | 2 | none | 6 |

**Table S1. 3D-MINFLUX scan parameters**

**Movie S1**. jGCamp8 calcium imaging of PIEZO1 clusters

**Movie S2.** 3D MINFLUX example with raw localizations of a GFP-labelled pit-shape PIEZO1 cluster

**Movie S3**. 3D MINFLUX example with raw localizations of a GFP-labelled spherical PIEZO1 cluster

**Movie S4.** 3D MINFLUX example with raw localizations and surface fit of a GFP-labelled pit-shape PIEZO1 cluster in control condition

**Movie S5.** 3D MINFLUX example with raw localizations and surface fit of a GFP-labelled pit-shape PIEZO1 cluster in hypo-osmotic condition

**Movie S6.** 3D MINFLUX example #1 of an ALFA-labelled and PIEZO1 expressing small neurite

**Movie S7.** 3D MINFLUX example #2 of an ALFA-labelled and PIEZO1 expressing small neurite
